# Neuronal networks in the dorsal hippocampus causally regulate rescue behavior in mice

**DOI:** 10.1101/2025.04.08.647754

**Authors:** Moisés dos Santos Corrêa, Anna Agafonova, Ada Braun, Charitha Omprakash, Esmeralda Tafani, Melissa Lowitsch, Leon Marquardt, Pavol Bauer, Anne Albrecht, Sanja Mikulovic

## Abstract

Prosocial behaviors, such as rescuing individuals in need, are crucial for social cohesion across species. While key brain regions involved in rescue behavior have been identified, the underlying neural mechanisms remain unclear. The hippocampus (HPC), known for its role in memory and spatial navigation, also contributes to emotional and social processing. However, its specific involvement in prosocial behavior is not well understood. Here, we investigate the causal role of the HPC in learning and executing rescue behavior in mice. Using chemogenetics, we show that the dorsal HPC (dHPC), but not the ventral HPC (vHPC), is essential for acquiring rescue behavior. Calcium imaging of the dHPC reveals network consolidation during successful rescues, with distinct synchronized ensembles and activity patterns linked to liberations of an individual in need. These findings establish a novel role for the dHPC in prosocial behavior, providing insights into the neural mechanisms underlying empathy-driven actions.

## INTRODUCTION

Empathy serves as a fundamental mechanism that binds social groups in humans and other animals by promoting prosocial behaviors^1,2^. While the presence of empathy and prosocial behaviors in non-human animals, including rodents, was once debated, it is now widely recognized^1–4^. Empathy is broadly defined as an involuntary emotional response that mirrors one’s perception of the emotional state of another individual^5^. Prosocial behaviors encompass any social actions aimed at alleviating distress and enhancing the well-being of others. Such behaviors have been observed across various animal species, including rescue behavior in ants^6^, birds^7,8^, dolphins^9^ and wild boars^10^, as well as helping behavior in elephants and non-human primates^11–13^. Prosocial behaviors are proposed to play a crucial role in the evolutionary success of social species by facilitating caregiving^14^, providing protection against predators, and ultimately enhancing genetic fitness through kin selection and inclusive fitness^12^.

While the first evidence of prosociality in rodents dates back to the 1950s and 1960s^15,16^, substantial progress in understanding the mechanisms behind prosocial behavior has been made over the past decade. Several robust behavioral paradigms have been developed to study different forms of prosociality, such as rescuing a trapped conspecific^17–19^, reward provision^20–22^, harm aversion^23–25^, and more recently pain alleviation and reviving-like behavior^26,27^. While most studies have focused on rats, mice—despite being less sociable^28^—were recently shown to also display helping behavior^29,30^. Although the extent to which prosocial behaviors in rodents are driven by underlying empathic motivation remains controversial^31–34^, it is widely accepted that emotional contagion contributes to modulating these behaviors^35,36^. Most mice exhibit emotional contagion, cognitive flexibility, and risk-taking behavior when learning to open a previously unfamiliar door to rescue companions^29,30^.

Recent studies have begun to provide key insights into the neurobiology of helping behavior. Several brain regions, similar to those identified as empathy-related hubs in humans^37–39^, have emerged as central modulators of helping. For example, it was shown that oxytocin silencing in the anterior cingulate cortex (ACC)^40^ and inhibition of the anterior insula^41^ reduced the latency for door opening to release trapped rats after a training period. ACC to nucleus accumbens (NAc) projections were shown to be specifically active while providing help to ingroup members^42^. Inhibiting the ACC reduced avoidance of a lever linked to harming another rat^23^ and impaired social licking of injured conspecific^27^ in mice. Although the HPC has been implicated in helping behavior in rodents^42^ and in the intention to help in humans^43–47^, no study has yet examined which specific aspects of helping behavior are encoded in the HPC. While the HPC is known for its role in memory formation^48^, it is not a uniform structure^49,50^. Functional differentiation along the dorsoventral hippocampal axis was first highlighted about 25 years ago, yet most research has focused on the dHPC, particularly for its role in cognitive and spatial processing^51^. Over the past decade, significant progress has been made in understanding the role of the vHPC, which has been shown to regulate emotion-related behaviors, including anxiety, stress, feeding, and social behavior^52^. However, the functional differentiation between the dHPC and vHPC is less discrete than previously assumed. Recent studies^43,45,53^ suggest that the dHPC is also involved in complex behaviors that include social and emotional components, indicating broader functional roles along the dorsoventral hippocampal axis.

Building on the robust establishment of a rescue paradigm in mice and the finding that emotional arousal of an individual in need enhances rescue behavior, we investigated a possible role of the HPC in regulating prosocial actions. Using a chemogenetic approach, we demonstrated that the dHPC, but not the vHPC, is causally involved in controlling the learning of rescue behavior. Furthermore, calcium imaging with miniscopes revealed a more engaged neuronal network over successful trials and distinct cell ensembles in the dHPC specifically associated with liberation behavior. C-Fos functional coactivity analysis comparing helper mice to dHPC-inhibited non-helper mice revealed a disruption in the co-activation of cognitive and emotional neural hubs following silencing of the dHPC. In summary, this study highlights the critical role of dHPC circuits in rescue behavior learning and paves the way for future research into their involvement in disorders affecting prosociality.

## RESULTS

### Emotional arousal of the victim promotes shorter liberation latencies

Negative emotional arousal in a trapped individual has previously been associated with shorter latency to help in rats^17,19,54^ and prairie voles^18^. To investigate if the same effect would be true to mice, we adapted a helping behavior task (HBT), previously described by Sato and colleagues^19^. In this task, we separated a pair of familiar mice into two compartments. One compartment contained the observer mouse, which had the opportunity to open an acrylic door, revealing a hole through which it could free the other mouse—the ‘victim’—trapped in the adjacent compartment. The trapped compartment was either empty (social separation group, SS, Figure S1A) or filled with cold water (cold water, CW, Figure S1B) to induce an emotionally aroused state of negative valence in the victim. As the observer explored its compartment, it could interact with the door and associate its opening with the liberation of the victim (Figure S1C). If the observer did not free the victim within the 15-minute session, the experimenter opened the door and guided the victim to the safe compartment. Over six days of training (Figure 1A) mice in the CW group had a higher probability of liberation of the victim (Figure 1B-C) and decreased the latency for emitting this behavior (Figure 1D). Comparing the learning of both groups, they showed similar learning rates over 6 days of training (Figure 1E). Observer mice were tested on day 7 with a different, non-familiar victim (Stranger test) and performed similarly well as during the last days of training (Figure 1D-E). Although our definition of rescuing relied primarily on the victim’s behavior (crossing the gap with its head), this action could only occur if the observer first opened the door. Thus, we interpret reduced latency over training sessions as evidence of learned rescue behavior—with the helper and victim adopting distinct but complementary roles in the task.

**Figure 1.**
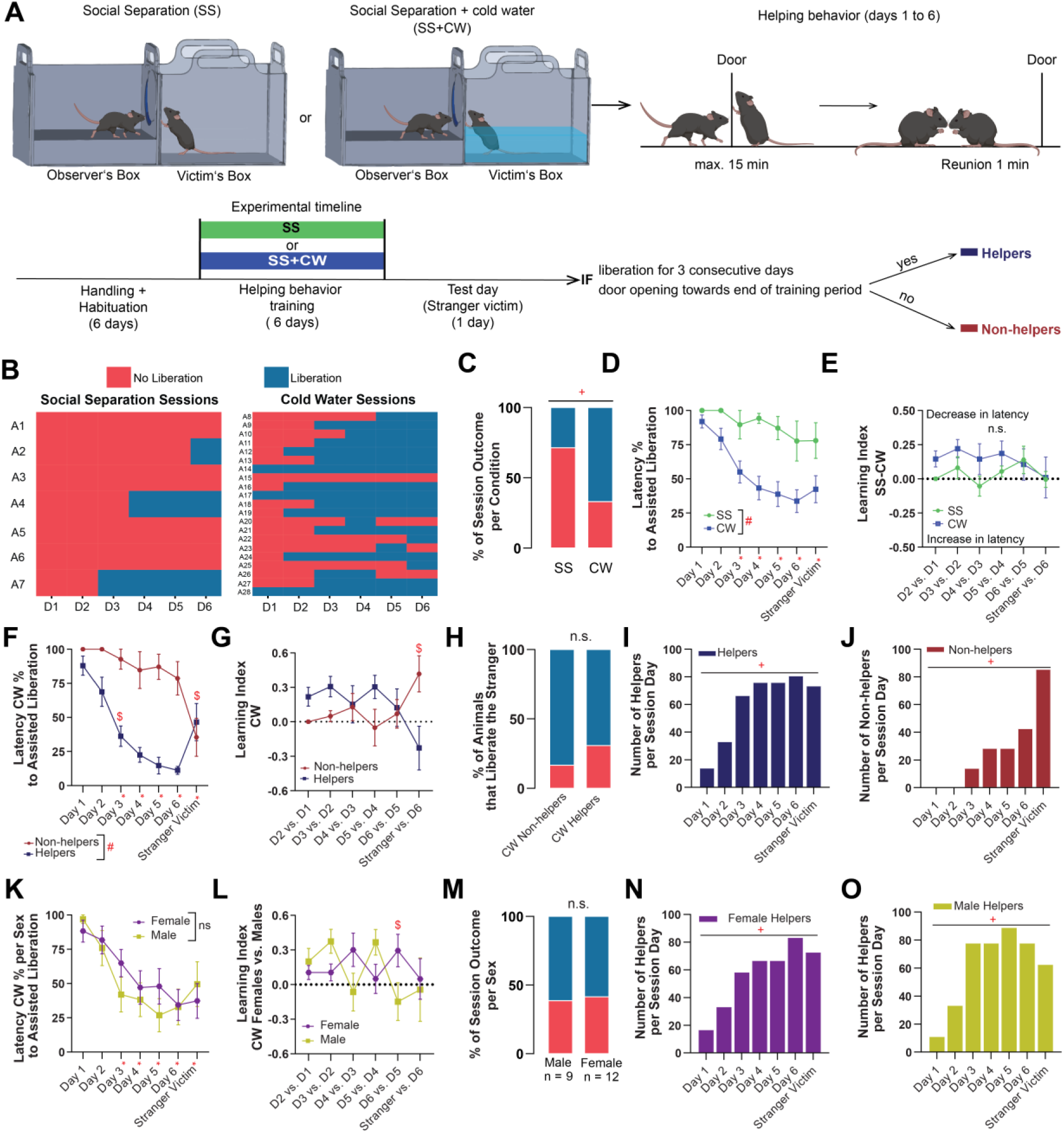
Emotional arousal of the victim promotes shorter liberation latencies in non-sex specific manner. (A-E) Validation of the Helping Behavior task using different levels of aversive conditions: Social Separation (SS) and additional Cold Water exposure (CW). (A) Schematic of experimental design, behavioral chambers, and training\testing sessions. (B) Matrices representing the session outcome for each pair of helper-victim mice from either the Social Separation (SS) only or Social Separation plus Cold Water (CW) groups, in red “no liberation” and in blue “liberation” (n = 7 in the SS group and 28 in the CW one). (C) Percentage of sessions with the session outcome “liberation” or “no liberation” in each condition (Chi-square with continuity correction: SS vs. CW, Χ²(1) = 20.6, p < 0.001). (D) Latency percentages for successful door opening and victim mouse liberation (Linear Mixed Model Omnibus test for Condition: F(1,24) = 12.26, p = 0.003, for Session: F(6,143.1) = 5.83, p < 0.001, interaction Session x Condition not significant; difference factor coding for Session to compare with all previous sessions and simple factor coding for Condition to compare groups). (E) Learning index calculated from each session’s latency value subtracted from its predecessor (LMM Omnibus test for Condition, Session and interaction, Condition x Session not significant). (F-J) Behavioral results comparing Helper and Non-helper mice within the CW condition (n = 14 Helpers and 7 Non-helpers). (F) Latency percentages for successful door opening and victim mouse liberation (LMM Omnibus test for Session: F(6,101.2) = 9.46, p < 0.001, for Outcome: F(1,17.1) = 51.36, p < 0.001; for interaction Outcome x Session day: F(6,101.2) = 5.43, p < 0.001; difference factor coding for Session to compare with all previous sessions and simple factor coding for Outcome to compare groups). (G) Learning index calculated from each session’s latency value with its predecessor (LMM Omnibus test for interaction Outcome x Session day: F(5,101) = 2.650, p = 0.027, factors Session and Outcome not significant). (H) Percentage of mice that liberate the non-familiar victim mouse during the Stranger test in the Helper or Non-helper groups (Chi-square: H vs. NH, Χ(1) = 0.4211, p = 0.5164). (I) Number of liberations per session day within the Helper group in the CW condition (Chi-square test for trend: Χ²(1) = 25.11, p < 0.0001). (J) Number of liberations per session day within the Non-helper group in the CW condition (Chi-square test for trend: Χ²(1) = 15.63, p < 0.0001). (K-O) Behavioral results comparing Females and Males within the CW condition. (K) Latency percentages for successful door opening and victim mouse liberation (LMM Omnibus test for Session: F(6,101.1) = 9.488, p < 0.001, factor Sex and interaction Session x Sex not significant. (L) Learning index calculated from each session’s latency value with its predecessor (LMM Omnibus test for interaction Condition x Session day: F(5,101) = 2.316, p = 0.049, factors Session and Sex not significant). (M) Percentage of sessions with the session outcome Helper or Non-helper in each condition when comparing sex differences within the CW condition (Chi-square with continuity correction: H vs. NH, Χ²(1) = 0.0196, p = 0.889). (N) Number of liberations per session day within the Helper group for female mice in the CW condition (Chi-square test for trend: Χ(1) = 13.32, p = 0.0003). (O) Number of liberations per session day within the Helper group for male mice in the CW condition (Chi-square test for trend: Χ(1) = 9.053, p = 0.0026). All data are presented as mean ± SEM. Outliers were identified and removed using the ROUT method. n.s. = non-significant, *p < 0.05 when comparing with all previous session days, #p<0.05 when comparing categorical groups, $p<0.05 when comparing interaction between previous sessions and groups, +p<0.05 in contingency tests.

As the CW condition proved to be a robust paradigm for investigating rescue behavior, we focused our subsequent analyses on this condition. To explore further how individual variability influences the learning of helping behavior, pairs of the CW group were divided the into two categories: helpers—those that demonstrated at least three consecutive rescue sessions towards the end of the training period—and non-helpers (Video S1). Helpers decreased the latency for liberating the victim over the training days and had a higher learning index compared to their conspecifics in the non-helper group. Surprisingly, both groups performed similarly well during the Stranger test (Figure 1F-H), albeit with contrasting learning index trends (Figure 1G). Observer mice in the non-helper group performed significantly better during the Stranger test relative to their late training performance (Figure 1I-J), indicating the importance of the victim identity for successful rescue behavior. Additionally, we examined sex differences in behavior of mice from the CW group. Female and male mice showed similar performances over training days and during the Stranger test (Figure 1K,M-O). However, differences in their learning indices suggest potential sex-specific variations in the learning rates over days (Figure 1L-O).

### Silencing of the dorsal but not ventral hippocampus impairs of helping behavior learning

Given evidence from human studies linking hippocampal lesions with impaired prosociality^44,55^ as well as the established functional diversity along the dorsoventral hippocampal axis^50^,we next focused on the role of the dHPC and vHPC in learning the rescue task (Figure 2A-B,M). We addressed their roles by injecting bilaterally adeno-associated virus (AAV) to express the inhibitory hM4Di receptor (iDREADD) in either the dHPC or vHPC of observer mice. Helping behavior training began approximately 3–4 weeks after viral injection. 30 minutes before each training session, observer mice received an intraperitoneal injection of the iDREADD agonist clozapine-N-oxide (CNO) or vehicle solution. To further examine the temporal dynamics of the HPC role in learning rescue behavior, we treated a third group of observer mice with a vehicle solution for five training days, but administered CNO only on the final day of training. Silencing the dHPC from the first training session onward (*CNO-all-days* group) dramatically reduced the expression of helping behavior in observer mice (Figure 2C-D). These animals also performed the task significantly worse across all training days compared to the *vehicle* or *CNO-last day* groups (Figure 2E-F). To determine whether CNO treatment affected other behavioral features, we used an automated behavioral tracking approach in combination with manual annotations (for details see Methods) to analyze locomotion, door interaction by observer mice, and social investigation between observer and victim mice after liberation. Similar to the vehicle group, CNO-all-days mice exhibited no preference for either the door or the opposite side of the safe chamber (Figure 2G). However, they displayed longer running bouts (Figure 2H), fewer door interaction bouts (Figure 2I), and reduced time investigating the area near the door prior to victim liberation (Figure 2J). These findings suggest a diminished motivation to rescue the conspecific in distress following the dHPC inhibition, despite maintaining the ability to interact with the door. There were no differences between *CNO-last day* and *vehicle* treated mice in any of these measurements. After victim liberation, *CNO-all days* mice spent a similar amount of time performing social investigation of the victim (Figure 2K) and had an equivalent rate of door interactions after liberation compared to the other two groups of mice (Figure 2L). These results demonstrate that dHPC inhibition causally regulates rescue behavior while leaving social memory intact. Although the vHPC has been recently associated with affective empathy^36,53,56^,it is not known whether vHPC engagement is necessary for learning prosocial behaviors. We applied a similar approach as in the dHPC silencing experiment, and targeted the vHPC during the 6 days of training and an additional Stranger test. Silencing the vHPC since the first day had no effect on the number of helpers in the observer mice (Figure 2N), the latency to liberate the victim mouse (Figure 2O), or the learning index, compared to the other experimental groups (Figure 2P). Analysis of other behavioral variables revealed no significant differences between groups, except that the vHPC *CNO-all-days* group spent significantly more time investigating the familiar victim mouse (Figure 2U) and consequently exhibited a lower rate of door interactions following the liberation event compared to the other groups (Figure 2Q–V). The dissociation between dHPC or vHPC silenced groups suggest a differential engagement of these regions to events before or after liberation, respectively.

**Figure 2.**
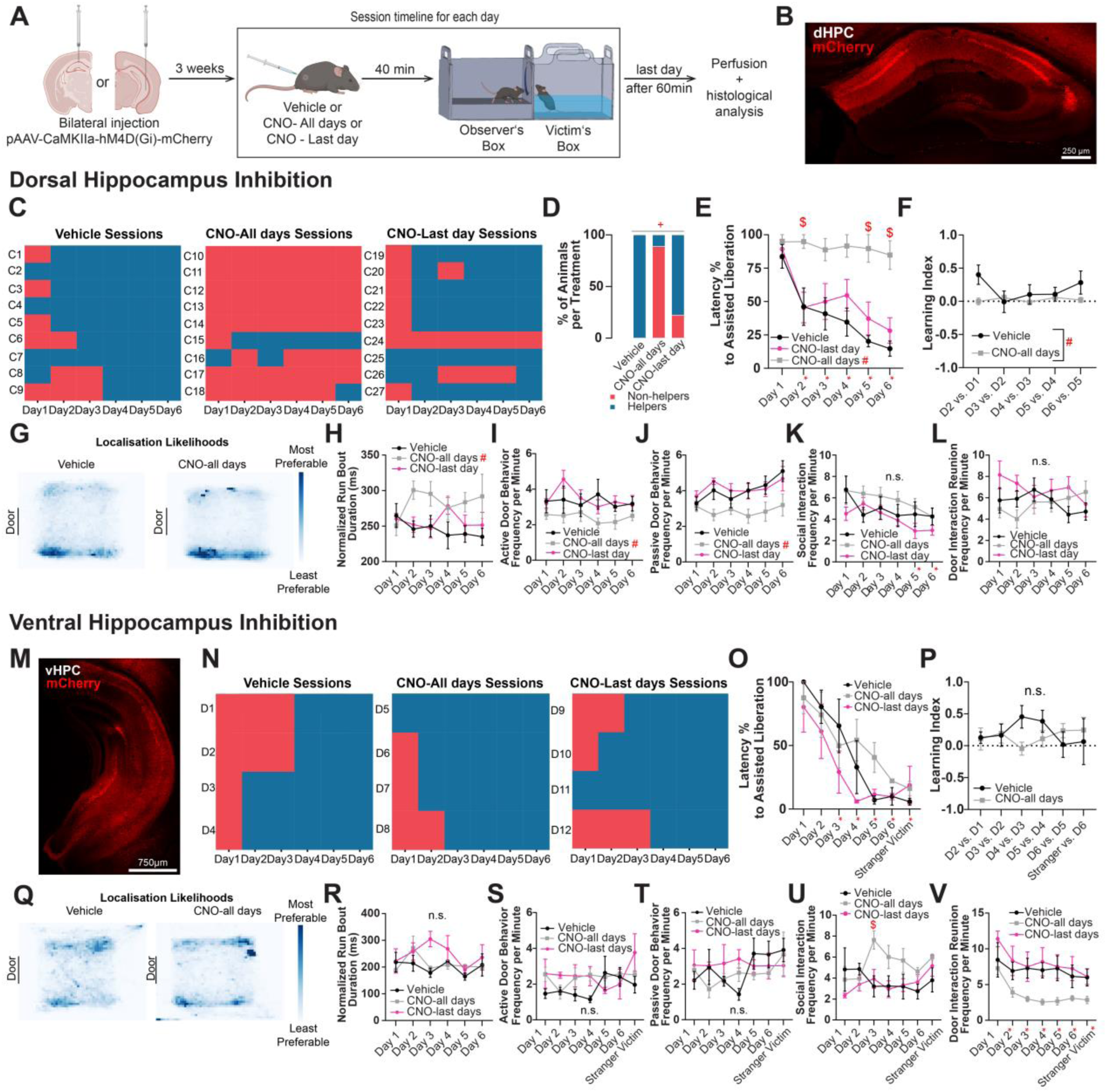
Silencing of the dorsal but not ventral hippocampus impairs helping behavior learning. (A) Schematic of infusion target sites and experimental design. (B-L) Behavioral results from the inhibition of the dHPC. (B) Representative image of virus spread (pAAV5-CaMKIIa-hM4D(Gi)-mCherry) in dHPC. (C) Matrices representing the session outcome for each pair of observer-victim mice from either the group of animals treated with *Vehicle* (Veh) before each training session, with CNO (*CNO-all days*) before each training session, or with CNO (*CNO-last day*) only before the last session, in red “no liberation” and in blue “liberation” (n = 9 in the Veh group, 9 in the *CNO-all days*, and 9 in the *CNO-last day*). (D) Percentage of sessions with the session outcome “liberation” or “no liberation” in each treatment (Chi-square: Χ(2) = 16.52, p = 0.0003). (E) Latency percentages for successful door opening and victim mouse liberation (Linear Mixed Model Omnibus test for factor Session: F(5,120.0) = 15.06, p < 0.001, for factor Treatment: F(2,24) = 13.24, p < 0.001, and for interaction Treatment x Session: F(10,120.0) = 2.87, p = 0.003; difference factor coding for Session and repeated factor coding for Treatment). (F) Learning index calculated from each session’s latency value subtracted from its predecessor (RM two-way ANOVA test for Treatment: F(1,16) = 9.93, p = 0.006; Holm-Sidak’s multiple comparison test). (G) Heatmap of the mean localization likelihoods of the observer mice for the Veh or *CNO-all days* group, indicating no change in preference for a specific location of the experimental chamber. (H) Normalized sum of running bout durations of the observer mice (LMM Omnibus test for Treatment: F(2,24.3) = 5.52, p = 0.011, for Session and interaction Session x Treatment not significant). (I) Frequency of active door interaction behavior bouts per minute of the observer mice (LMM Omnibus test for Treatment: F(2,24.0) = 5.44, p = 0.011, for Session and interaction Session x Treatment not significant). (J) Frequency of explorative behaviors bouts of the observer mice occurring near the area of the door, but with no direct door interaction (LMM Omnibus test for Treatment: F(2,24.0) = 7.94, p = 0.002, for Session and interaction Session x Treatment not significant). (K) Frequency of social exploration behaviors between the observer and victim mice per minute after liberation (LMM Omnibus test for Session: F(5,120.0) = 3.24, p = 0.009, for Treatment and interaction Session x Treatment not significant). (L) Frequency of door interaction behavior bouts of the observer mice per minute (LMM Omnibus test for Session, Treatment and interaction Session x Treatment not significant). (M-V) Behavioral results from the inhibition of the ventral hippocampus (vHPC). Here, the group *CNO-last days* was treated with CNO in both Day 6 and the Stranger Victim test. (M) Representative image of virus spread in vHPC. (N) Matrices representing the session outcome for each pair of helper-victim mice from either the Veh, *CNO-all days*, or *CNO-last days*, in red “no liberation” and in blue “liberation” (n = 4 in the Veh group, 4 in the *CNO-all days*, and 4 in the *CNO-last days*). (O) Latency percentages for successful door opening and victim mouse liberation (LMM Omnibus test for Session: F(6,54) = 22.03, p < 0.001; test for Treatment and interaction Treatment x Session not significant). (P) Learning index calculated from each session’s latency value subtracted from its predecessor (LMM Omnibus test for Session, Treatment and interaction Treatment x Session not significant). (Q) Heatmap of the mean localization likelihoods of the observer mice for the Veh or *CNO-all days* group, indicating no change in preference for a specific location of the experimental chamber. (R) Normalized sum of running bout durations of the observer mice (LMM Omnibus test for Session, Treatment and interaction Treatment x Session not significant). (S) Frequency of active door interaction behavior bouts per minute of the observer mice (LMM Omnibus test for Session, Treatment and interaction Treatment x Session not significant). (T) Frequency of explorative behaviors bouts of the observer mice occurring near the area of the door, but with no direct door interaction (LMM Omnibus test for Session, Treatment and interaction Treatment x Session not significant). (U) Frequency of social exploration behaviors per minute between observer and victim mice after liberation (LMM Omnibus test for interaction Treatment x Session: F(12,53.05) = 2.28, p = 0.020, for factors Session and Treatment not significant). (V) Frequency of door interaction behavior bouts per minute performed by the observer mouse after victim liberation (LMM Omnibus test for Session: F(6,52.07) = 8.02, p < 0.001, for Treatment and interaction Session x Treatment not significant). All data are presented as mean ± SEM. n.s. = non-significant, *p < 0.05 when comparing with all previous session days, #p<0.05 when comparing categorical groups, $p<0.05 when comparing interaction between previous sessions and groups, +p<0.05 in contingency tests.

### Learning to help is associated with increased engagement of the dorsal CA1 neurons and their functional coactivity over days

Given our above-described findings showing the critical role of the dHPC in rescue behavior, we next investigated whether changes in the dorsal CA1 (dCA1) network correlate with the number of rescue trials over time, reflecting hippocampus-dependent learning. We chose the dCA1 since it encodes multi-dimensional representation of episodic spatial context^57^, form cell assemblies that encode features of individual states^58^ and supports retrieval of sequences of episodic events inside a spatial context^59^. We reasoned that these features are essential for rescuing an individual in need within an environment.

To monitor neuronal activity in the dCA1, we injected the calcium indicator GCaMP8m and implanted a GRIN lens in the same region. We then applied functional connectivity graph analysis to examine cell ensemble dynamics during rescue behavior (Figure 3A–C). Miniscope-implanted mice were first trained in the HBT under freely moving conditions (Figure S1D, Video S2) and subsequently tested on separate days either with a stranger victim or in a novel spatial context featuring distinct spatial cues (Recent CtxB test, Figure S1E). Twenty-eight days later, we conducted additional tests in both the novel (Remote CtxB test) and original training (Remote HBT test) contexts with the familiar victim, while imaging continuously throughout all behavioral sessions. Finally, we exposed miniscope observer mice to victimless control test days. Here, the victim was exchanged with either a toy mouse, a pool of cold water, or an empty chamber (Figure S1F-H). As reported above, most observer mice learned to liberate the trapped victim faster over days, but during the test days, they took in average longer (Figure 3D-E). Following the extraction of spatial footprints and time series of detected neurons, we observed a progressive decrease in the percentage of identified cells per observer animal across training days. However, the identified neurons remained stable during test days, except for the Remote CtxB (Figure 3F). Furthermore, the percentage of identified neurons during the training days was positively correlated with the latency to liberate the trapped victim (Figure 3G). Since session length in our dataset is contingent on mouse performance (i.e., quicker rescues lead to shorter calcium recordings), we hypothesized that longer sessions may enable the detection of a greater number of neuronal components, or alternatively, that faster rescues involve the recruitment of fewer neurons. To test for a possible facilitatory effect on the number of detected components and to control subsequent analysis steps, we generated a synthetic dataset of randomly firing components and processed the *in silico* videos using the same pipeline as the *real* dataset (Figure S2). The *in silico* dataset replicates all the major features of the real data — including spatial footprints, session duration (as indicated by helping latency), and neuronal signal profiles (Figure S3A–C) — but with randomly timed firing and with higher signal-to-noise ratios (Figure S3D). Unlike the *real* data, the number of components in the *in silico* data remained stable over time and was not correlated with the latency (Figure S3E–F). Thus, we concluded that as animals progressively learn to perform rescue behaviors across multiple trials, the dCA1 network becomes more efficient, requiring the activation of fewer cells. This claim is further supported by an increase of the calcium event rates over rescue trials (Figure 3H-J), which is not the case for the *in silico* dataset (Figure S3G). Victimless control days resulted in lower event rate compared to when mice had 5 or more successful rescuing trials (Fig. S5C).

**Figure 3.**
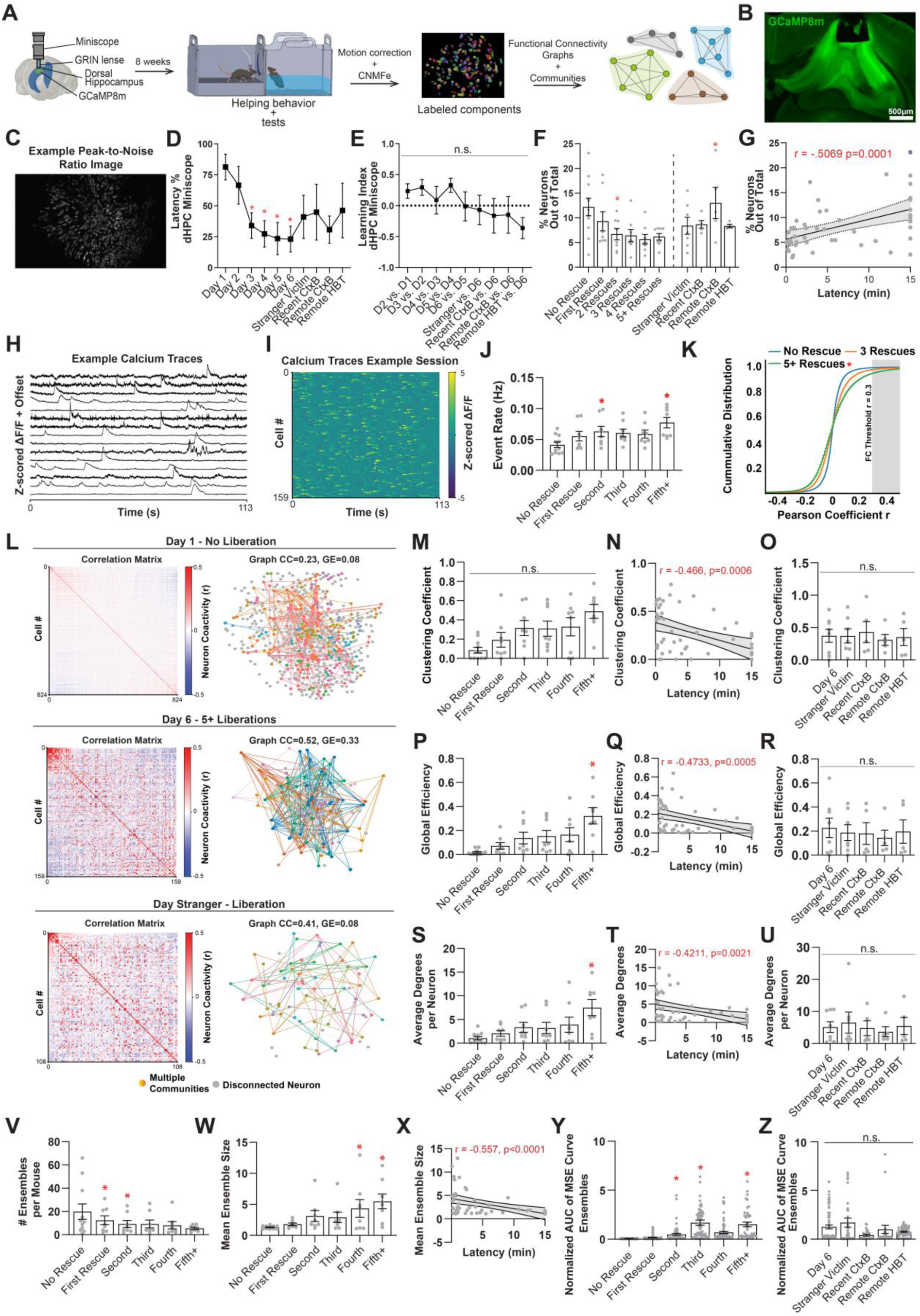
Learning to help is associated with increased engagement of CA1 neurons and their functional coactivity over days. (A) Schematic of miniscope calcium imaging of dorsal hippocampus CA1 neurons of freely moving mice trained on the helping behavior task. After cell identification and extraction of ΔF/F signals, functional connectivity analysis was performed to identify pairs of cells that had reasonably correlated calcium events. Graph analysis was then performed to verify the clustering coefficient and global efficiency of the network across days of training and testing. (B-I) Behavioral and calcium imaging components evaluation results of the dorsal CA1. (B) Representative histology image of virus spread (pGP_AAV-syn_jGCaMP8m-WPRE) and lens placement above the CA1 layer of the dHPC. (C) Representative frame taken from one the acquired endoscopic videos during the helping behavior training. (D) Latency percentages for successful door opening and victim mouse liberation (LMM Omnibus test Session: F(9,54.2) = 3.10, p = 0.005; difference factor coding for Session). (E) Learning index calculated from each session’s latency value subtracted from its predecessor (RM two-way ANOVA test for Session not significant). (F) Percentage of cells found in given mouse from the maximum number of cells present in this mouse across all sessions grouped across days of helping starting from „No Rescue” to “5 or more rescues” and subsequent test sessions (LMM Omnibus test for interaction Day of Helping: F(9,62.0) = 2.60, p = 0.013, difference factor coding for Day of Helping). (G) Pearson correlation between cell percentage and latency for helping. (H) Representative z-scored ΔF/F signals from 15 cells acquired from one of the mice during one of the training sessions. (I) Representative heatmap from all cells acquired from one the mice during in one of the training sessions (J) Event rate (Hz) per animal across days of helping (LMM Omnibus test for interaction Day of Helping: F(5,37.2) = 4.14, p = 0.004, difference factor coding for Day of Helping). (K) Cumulative distributions of Pearson’s coefficients per day of helping, gray area indicates values r ≥ 0.3: (Sample sizes: No rescue N=275,156; First rescue N = 60,738; 5+ rescues N = 61,320 correlations). Bootstrapped two-sample Kolmogorov–Smirnov tests revealed significant differences between No Rescue (blue) and Day 5+ (green) (KS = 0.197 ± 0.057 CI, p = 0.048, corrected) (L-U) Functional coactivity analysis for all animals, including one non-helper. Pairwise Pearson correlations for all approved cells from each mouse and session were compared to their counterparts from same animal and session. Graphs were then constructed using all correlations which coefficients are over a threshold of +0.30. Neurons were grouped into different ensembles according to the above-threshold correlations using the Leiden algorithm. Graph properties were then evaluated by measuring the clustering coefficient, global efficiency and average degree per neuron of the network across days of helping. (L) Representative correlation matrices (left) and corresponding graphs (right) for one animal at the day 1 (upper panels) and day 6 (middle panels) of training, and on the stranger victim test day (bottom panels). In the representative matrices, positive Pearson coefficients are color coded in red whereas negative coefficients are coded in blue. In the representative graphs, each dot is a neuron arranged approximately at the real location in the endoscopic video and each line represents a correlation which coefficient is over +0.30. Gray circles represent disconnected neurons whereas other colored circles represent different unique ensembles. CC means clustering coefficient and GE means global efficiency. (M) Clustering coefficient across days of rescue: (LMM Omnibus test for factor Day of Helping: F(5,37.5) = 2.13, p = 0.083). (N) Pearson correlation between clustering coefficient and latency for helping. (O) Clustering coefficients across test sessions: (LMM Omnibus test for interaction Session not significant). (P) Global efficiency of the network across days of helping (LMM Omnibus test for Day of Helping: F(5,37.8) = 3.69, p = 0.008). (Q) Pearson correlation between global efficiency and latency for helping. (R) Global efficiency across test sessions: (LMM Omnibus test for interaction Session not significant). (S) Average degrees per neuron across days of rescue: (LMM Omnibus test for Day of Helping: F(5,37.9) = 2.59, p = 0.041). (T) Pearson correlation between average degrees and latency for helping. (U) Average degrees per neuron across test sessions: (LMM Omnibus test for interaction Session: not significant). (V-Y) Metrics related to ensembles of at least 2 neurons, extracted using the Leiden algorithm for community detection. (V) Number of ensembles per mouse across days of rescue: (LMM Omnibus test for Days of Helping: F(5,38.1) = 3.41, p = 0.012). (W) Average ensemble size per mouse across days of rescue: (LMM Omnibus test for Days of Helping: F(5,38.4) = 2.75, p = 0.032). (X) Pearson correlation between mean ensemble size and latency for helping. (Y) Normalized multi-sample entropy (MSE) area under curve value per ensemble across days of rescue: (LMM Omnibus test for Days of Helping: F(5,558.0) = 42.2, p < 0.001). (Z) Normalized MSE area under curve value per ensemble across test sessions: (LMM Omnibus test for Days of Helping not significant). All data are presented as mean ± SEM. n.s. = non-significant, *p < 0.05 when comparing with all previous rescue trials. Outlier datapoints were removed using the ROUT test.

To further investigate changes in the dCA1 network during rescue trials, we analyzed functional coactivity using pairwise Pearson’s correlation coefficient (PCC) matrices. Recent studies used functional coactivity from the extracted time series of populations of neurons in the HPC, to investigate how ensembles of cells correlate their activities during learning tasks^61–63^. We observed that the *real* network became even more functionally coactive after five or more trials compared to three or zero successful trials (Figure 3K). This increase was absent in the *in silico* dataset (Figure S3H). To also evaluate the validity of our approach presented above, we next compared the distribution of PCC values in *real* versus *in silico* datasets (Figure S4A). We focused on the distribution tails—particularly the positive extreme, where neurons exhibit highly synchronized activity (Figure S4B). Notably, the *real* dataset contained a significantly greater number of highly correlated pairs compared to the *in silico* dataset (Figure S4C–D). To confirm that these high correlations were not driven by extrinsic factors, we examined the relationships between PCC and physical distance, signal-to-noise ratio (SNR), and spatial correlation (Figure S4E–G). This analysis showed no dependency between correlation and these extrinsic parameters, supporting the idea that high correlations reflect an intrinsic property of the real neuronal network.

Having observed a progressive increase in neuronal functional connectivity across training days, we next examined whether additional network properties also changed. Using identity correlation matrices for each session, we constructed binary, undirected graphs representing functional coactivity in dCA1 and applied the Leiden community detection algorithm to identify neuronal ensembles (Figure 3L). We then assessed network dynamics over the course of rescue training using established graph metrics: clustering coefficient (the degree to which nodes cluster), global efficiency (inverse of the shortest path length), and average degree per neuron (number of edges per node, reflecting node strength)^64,65^. We observed that the dCA1 network exhibited a progressive increase in clustering coefficient during later successful rescue trials (Figure 3M), a trend that was associated with shorter liberation latencies (Figure 3N). Notably, the clustering coefficient remained stable during test sessions (Figure 3O). Similar patterns were found for global efficiency and average degree per neuron, both of which increased across training trials (Figure 3P,S) and negatively correlated with rescue latency (Figure 3Q,T). These network metrics remained stable during test sessions, suggesting that once acquired, the neural representation of rescue behavior is maintained across varying social, spatial, and temporal contexts (Figure 3R,U). In contrast, sessions without a victim resulted in significantly lower clustering coefficient, global efficiency, and average degree compared to sessions following five or more successful rescues (Figure S5D–F). Importantly, while network visualizations of the *in silico* data showed no obvious changes over time (Figure S6C), the corresponding metrics remained consistently lower than in *real* data (Figure S6A,D,H) and showed different changes across rescue days (Figure S6B,E,I), further validating the specificity of our findings.

Next, to assess whether neuronal clustering leads to ensemble formation across trials, we analyzed community-related metrics within the dCA1 graphs. We observed that the number of ensembles (defined as at least 2 functionally coactive neurons) decreased over rescue trials (Figure 3V), but the average size of these emerging ensembles increased (Figure 3W). The mean ensemble size was inversely correlated with latency, indicating that better rescue performance recruits more functionally connected neurons (Figure 3X). The mean ensemble size on victimless control days was smaller during Toy, Pool, and Empty sessions compared to days when mice completed five or more successful rescue trials (Figure S5G). Comparison with the *in silico* data revealed that the number of ensembles over trials tends to increase over days, but that both *real* and *in silico* datasets exhibited similar ensemble sizes (Figure S6G). To further assess the variability of the ensemble calcium signals during learning, we applied multi-scale Sample Entropy (MSE; see Methods) to the average activity of each ensemble. After normalizing by latency times, we observed that MSE for these ensembles increased over rescue trials, with mice showing values close to 0 on No Rescue or First Rescue trials, while significantly higher values from the second rescue on (Figure 3Y). *In silico* data, on the other hand, maintained values close to 0 over all trials (Figure S6J,K). Interestingly, only one observer mouse from the Miniscope sample was categorized as non-helper according to the criteria described previously (Figure S7A). This mouse, compared to the helper observer mice during No Rescue sessions or after 5+ rescue trials, showed decreased mean event rate (Figure S7B). Functional coactivity for this mouse was comparable to the No Rescue sessions of the helper mice in all metrics analyzed (Figure S7C-E). Community analysis showed fewer ensembles of coactive neurons and increased ensemble MSE when compared to helper mice (Figure S7F-G). Reduced hippocampal engagement during HBT training may underlie the failure to successfully acquire rescue behavior. Together, these results indicate that the dCA1 network is structurally organized and dynamically shaped by experience during rescue behavior learning.

### Liberation and non-liberation neuronal ensembles in dCA1 show differential activity patterns over rescue behavior training days

Previous studies have shown that neurons in the dCA1 present internal generation of cell-ensemble sequences and these sequences reflect information about action planning and behavioral performance^60,61^. Additionally, functionally connected activity in dCA1 has been suggested to encode representation of the spatial context where a learned association occured^62^. Our previous results suggested that the rescuing paradigm may reflect respondent learning that is possibly dependent on spatial, social or emotional components (Figure 3D-F). This raises the question of whether the activity patterns of functionally connected neuronal ensembles systematically change across training days or test sessions, which can indicate learning-related plasticity. For this, we investigated the correlation of calcium traces (synchronization score), calcium activity as well as the MSE within each ensemble. Our results indicated that whole-session ensemble synchronization score did not change during training and test sessions (Figure 4A-B). We then defined the ‘liberation window’ around the liberation event. Liberation-related calcium activity (AUC of calcium peaks in the liberation window) exhibited non-significant changes across the training days when all ensembles were taken into account, except during ‘No Rescue’ sessions, where the activity was significantly reduced (Figure 4C–D). This suggests greater involvement of the dHPC in self-initiated liberation compared to experimenter-initiated release. Importantly, neuron synchronization within ensembles during the liberation time interval compared to the rest of the session increased with successful training trials (Figure 4E), but decreased during test sessions (Figure 4F), suggesting a dynamic, context-dependent reorganization of neuronal ensembles associated with rescue behavior.

**Figure 4.**
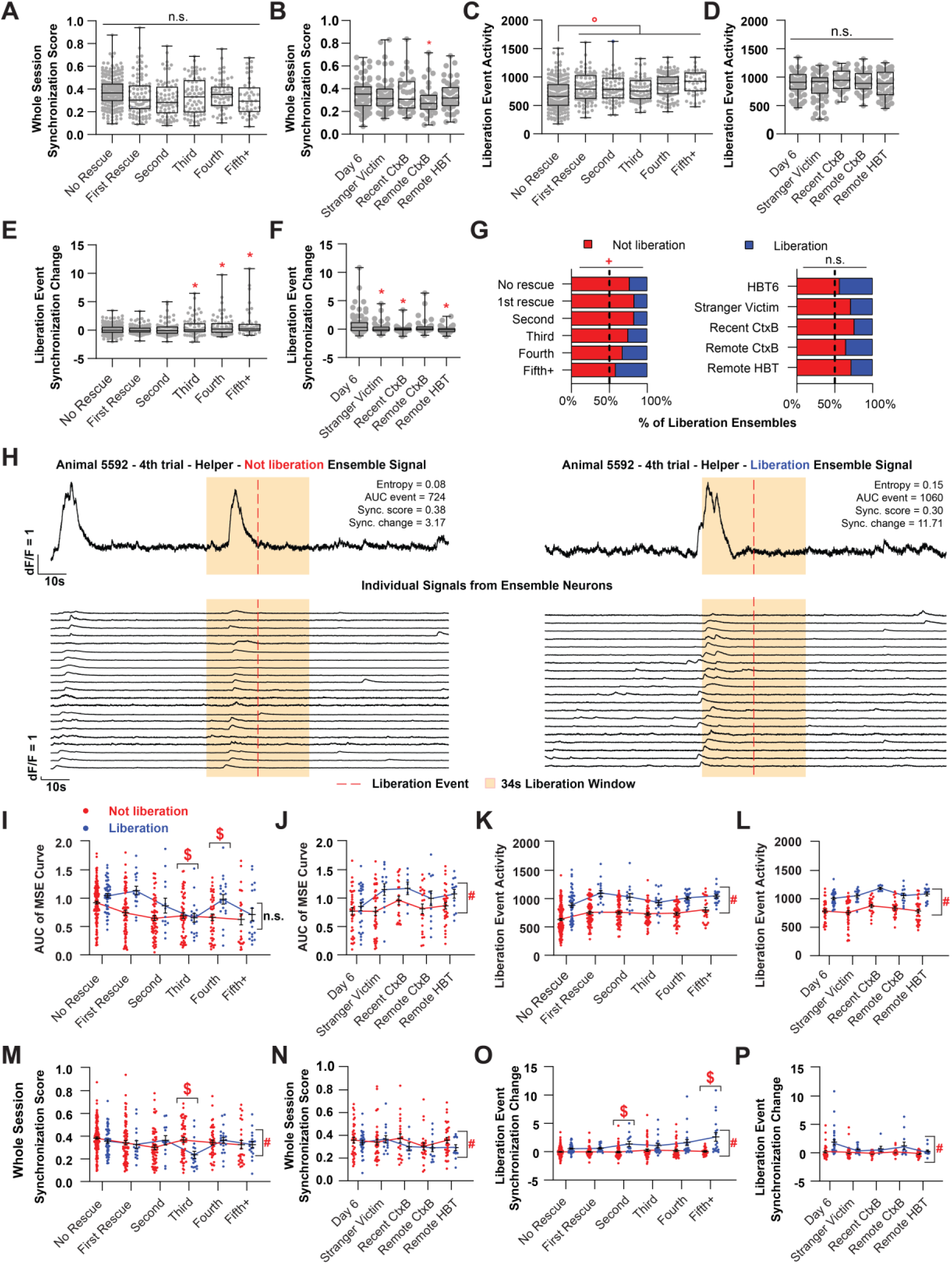
Liberation and non-liberation ensembles in dCA1 show differential neural activity patterns over helping behavior training days. (A-F) Calcium activity during liberation event and synchronization score results for the averaged signal of all ensembles of at least 2 neurons, for both training and test sessions. Ensembles were extracted from all animals and sessions, regardless of outcome (helper or no-helper). Calcium activity during liberation event was measured by detecting peaks inside the 34 seconds time interval around the victim head crossing and calculating the area under curve (AUC) for all detected events. Synchronization score is the averaged Pearson correlation coefficient for the whole signal of all neurons in the ensemble. Synchronization change is the ratio between the score calculated only for the 34s liberation window and the score for the rest of the session. (A) Whole session synchronization scores for all ensembles across rescue days: (LMM Omnibus test for factor Day of Helping not significant). (B) Synchronization scores across test sessions: (LMM Omnibus test Session: F(4,217) = 2.44, p = 0.048; simple factor coding for Session). (C) Liberation event activity as AUC of calcium events during the liberation window for all ensembles across rescue days: (LMM Omnibus test for factor Day of Helping: F(5,558) = 12.60, p < 0.001, helmert factor coding for Day of Helping comparison). (D) Liberation event activity across test sessions: (LMM Omnibus test Session not significant). (E) Ratio of synchronization change during liberation window for all ensembles across rescue days: (LMM Omnibus test for factor Day of Helping not significant). (F) Ratio of synchronization change across test sessions: (LMM Omnibus test Session: F(4,109) = 4.15, p = 0.004; simple factor coding for Session). (G) Percentage of non-liberation and liberation ensembles across rescue days (Chi-square: Χ(5) = 14.3, p = 0.014) or test sessions (Chi-square: Χ(4) = 6.57, p = 0.161). (H) Representative figures for a non-liberation ensemble (right side) and a liberation one (left). Both figures depict the averaged signal between all member neurons (above) and signals from each individual neuron in that ensemble (below). For a fair comparison, both ensembles belong to the same mouse, from the same training day. The shaded area represents the 34s-time window around the moment which the victim mouse crossed the head through the door, and the red dashed line represents this moment. The non-liberation ensemble shows two peaks, one outside the liberation window and another inside. The liberation ensemble shows only one peak, inside the liberation window. The signal from individual neurons show how correlated calcium traces were for both ensembles. The metrics for both ensembles are also described in the panel. (I-P) MSE, calcium activity and synchronization score results for liberation and non-liberation ensembles, for both training and test sessions. (I) Multi-Sample Entropy (MSE) AUC value comparing non- and liberation ensembles across days of rescue: (LMM Omnibus test for Day of Helping: F(5,550) = 8.01, p < 0.001, for interaction Day of Helping x Liberation: F(5,549) = 2.81, p = 0.016, for Liberation not significant; difference factor coding for Day of Helping, simple factor coding for Liberation). (J) MSE comparing non- and liberation ensembles across test sessions: (LMM Omnibus test Liberation: F(1,212) = 11.702, p < 0.001, for factor Session and interaction Session x Liberation not significant). (K) Liberation event activity for non- and liberation ensembles across rescue days: (LMM Omnibus test for Day of Helping: F(5,551) = 10.06, p < 0.001, for Liberation: F(1,552) = 112.89, p < 0.001, interaction Day of Helping x Liberation not significant; helmert factor coding for Day of Helping). (L) Liberation event activity comparing non- and liberation ensembles across test sessions: (LMM Omnibus test Liberation: F(1,212) = 129.95, p < 0.001, for Session and interaction Session x Liberation not significant). (M) Whole session synchronization scores comparing non-liberation and liberation ensembles across rescue days: (LMM Omnibus test for Liberation: F(1,556) = 4.60, p = 0.032, for interaction Day of Helping x Liberation: F(5,552) = 2.81, p = 0.016, for Day of Helping not significant). (N) Synchronization scores comparing non- and liberation ensembles across test sessions: (LMM Omnibus test for Session: F(4,215) = 2.45, p = 0.047, for Liberation: F(1,212) = 4.51, p = 0.035, for interaction Session x Liberation not significant). (O) Ratio of synchronization change during liberation window for non- and liberation ensembles across rescue days: (LMM Omnibus test for Day of Helping: F(5,532) = 7.21, p < 0.001, for Liberation: F(1,555) = 107.17, p < 0.001, for interaction Day of Helping x Liberation: F(5,553) = 5.96, p < 0.001; difference coding factor Day of Helping and simple factor coding for Liberation). (P) Ratio of synchronization change comparing non- and liberation ensembles across test sessions: (LMM Omnibus test for Session: F(4,193) = 3.93, p = 0.004, for Liberation: F(1,213) = 27.98, p < 0.001, for interaction Session x Liberation not significant). All data are presented as mean ± SEM. n.s. = non-significant, *p < 0.05 when comparing with all previous days of helping, #p<0.05 when comparing categorical groups, $p<0.05 when comparing interaction between previous sessions and groups, +p<0.05 in contingency tests, °p < 0.05 when comparing to next days of helping.

Next, we reliably classified neural ensembles as either ‘liberation’ or ‘non-liberation’ based on the probability of calcium events occurring during the liberation event versus outside this time window (Figure 4G-P). We then examined how synchronization scores, calcium activity, and MSE evolved across these ensembles over successive days. We found that the proportion of liberation ensembles increased over successful rescue trials and remained stable during test days (Figure 4G). By comparing the average neuronal activity patterns between ‘liberation’ and ‘non-liberation ensembles’, we found significant differences in MSE during the third and fourth rescuing trials compared to the first, second or no rescue (Figure 4I). Additionally, on the test days, ‘liberation ensembles’ exhibited higher MSE than ‘non-liberation’ ones (Figure 4J). Calcium activity during the liberation window was higher during training and testing sessions for ‘liberation ensembles’ (Figure 4 K-L). Throughout the session, synchronization scores were slightly higher in ‘liberation ensembles’ during both training and test sessions (Figure 4M-N). In contrast, the event synchronization change in liberation ensembles is significantly higher in later training trials when compared to non-liberation ones (Figure 4O), and this increase drives the effect seen in Figure 4 E. Together, these findings suggest that a subset of neuronal ensembles becomes increasingly responsive and selectively tuned to the liberation event over the course of rescue training. The persistence of certain activity patterns during test sessions, despite contextual changes, indicates that these ensembles may encode core features of the learned rescue behavior. Reduced synchronization during tes points to a degree of representational flexibility, potentially supporting adaptation to novel social, spatial, or temporal contexts.

### Silencing of the dHPC during rescue behavior training elicits changes in functional coactivity of brain-wide c-Fos expression

After identifying and characterizing the ‘liberation’ and ‘non-liberation’ ensembles, we next sought to investigate additional brain regions potentially influenced by the dCA1 encoding of rescuing behavior, as well as the impact of dHPC silencing on these regions. First, we performed immunostainings for the immediate early gene c-Fos in mice that were previously injected with AAV5-CaMKIIa-hM4D(Gi)-mCherry in the dHPC and either trained after being treated with Vehicle or CNO for all 6 sessions. As previously reported in Figure 2, most animals of the *CNO all-days* group failed to learn to rescue the victim during the training period. We pre-selected regions that have been previously associated with helping behavior, emotional learning or episodic memory, such as vHPC subregions (vDG, vCA3 and vCA1), the basolateral nucleus of the amygdala (BLA), septal subregions (medial and lateral septum, MS and LS), the anterior cingulate (ACC), the retrosplenial (RSC) cortices, and the subregions of the dHPC (dDG, dCA3, dCA2 and dCA1)^42,63^. Compared to the *vehicle* group, the *CNO all-days* animals showed increased c-Fos expression in the vCA3, vCA1, and the RSC (Figure 5B). We additionally observed tendency of increase of c-Fos-positive cells in the BLA of *CNO all-days* mice (Figure 5B). We next investigated functional coactivity (FC) of c-Fos expression between these regions and found that the *CNO all-days* group showed a differential engagement of emotional (BLA and vHPC) and cognitive (ACC, RSC and dHPC) nodes, with higher clustering compared to the *vehicle* group (Figure 5 C-D). Although these two ensembles were identified in both groups, the *CNO all-days* group showed higher clustering (Figure 5D). The *vehicle* group, on the other hand, showed more efficient and higher coactivity of hippocampal subregions, both in the dorsal and ventral portions (Figure 5D). We then calculated the difference between the groups across all pairwise correlations which indicated that the *CNO all-days* group displayed an increased coactivity between the dDG and dCA3 to the RSC, the dCA2 to the vDG and the BLA to the vCA1 (Figure 5E). Lastly, we performed a network analysis of centrality, closeness, clustering and degrees to identify central hubs among the investigated regions. For the *vehicle* group, we identified the dDG and vCA1 as central hubs of the more efficient network, whereas for the *CNO all-days* group the BLA was the only central hub that connected both highly clustered ensembles (Figure 5F). Overall, these findings suggest that silencing the dHPC increases the functional connectivity modularization of emotional and cognitive brain regions. This effect may result from removing an integrative hub from the network, which in turn negatively impacted the learning of rescue behavior.

**Figure 5.**
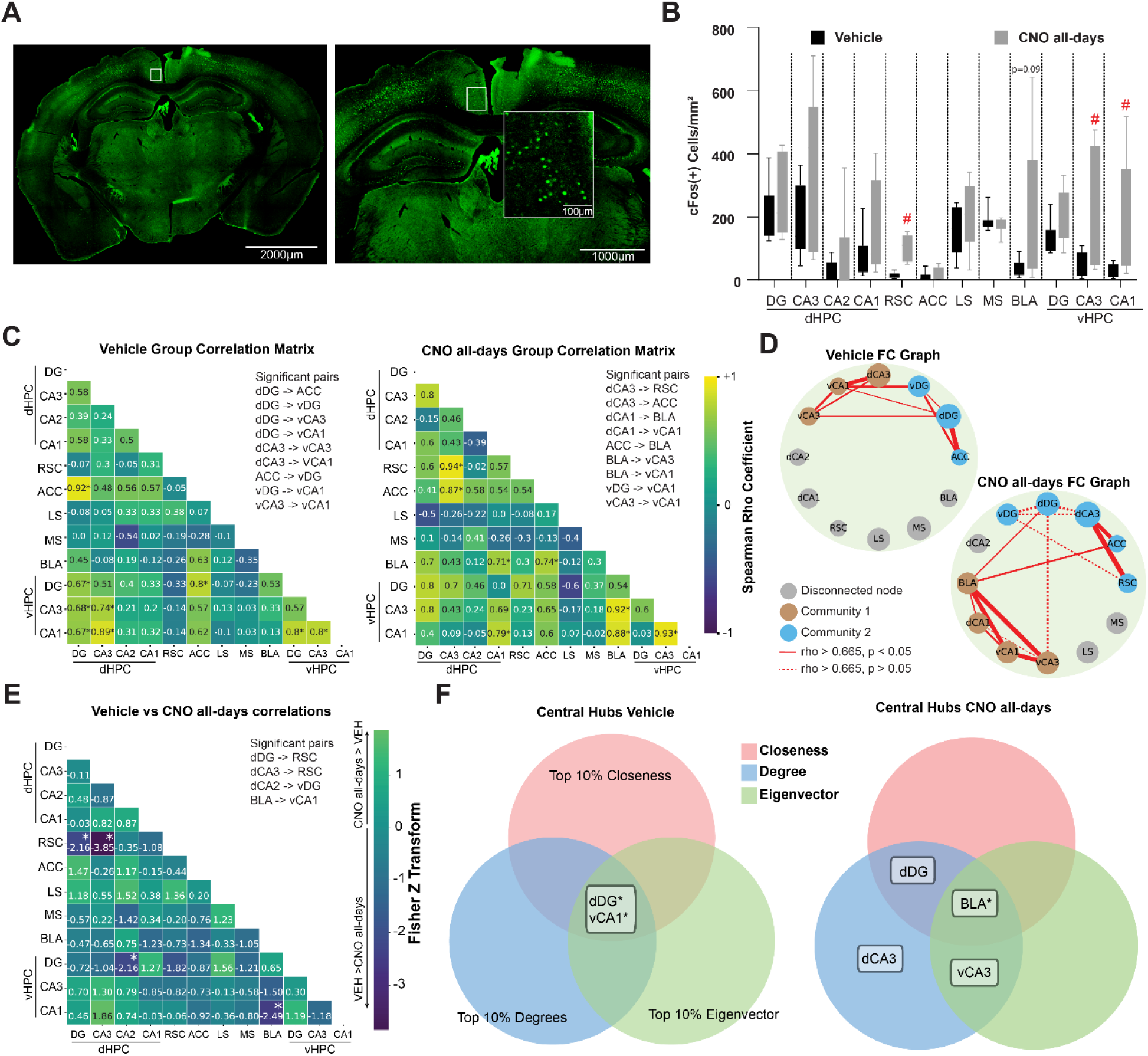
Silencing the dHPC during helping behavior training elicits changes in functional coactivity of c-Fos expression in different brain regions. (A) A representative image of c-Fos signal (green) sampled in the RSC. (B) Differences in c-Fos+ cells between the mice that were treated either with vehicle (Vehicle group) or CNO (CNO all-days group) before every training session in each of the investigated brain regions. Density of c-Fos+ cells per area across brain regions (Mann-Whitney tests between Vehicle and CNO all-days – RSC: U = 0, p < 0.001; vCA3: U = 14, p = 0.019; vCA1: U(57,114) = 12, p = 0.011). (C-D) Coactivity and graph analysis of c-Fos expression between all investigated brain regions in both groups. (C) Heatmap matrix representing all pairwise Spearman correlations between each investigated brain region of the Vehicle (left) or CNO all-days (right) groups. Rho coefficient values range +1 (yellow) to −1 (dark blue). Coefficient values and significance symbols are written in each bin of the matrix. (E) Graph representing all investigated brain regions from the Vehicle (top-left) or CNO all-days (bottom-right) groups, with the circles of different colors representing the nodes and the edges between them representing the correlation above the threshold of rho = 0.665. Circle sizes represent the normalized c-Fos+ density for that brain region. Circle color represents clusters identified via a Leiden algorithm. Edge thickness represents the normalized rho coefficient between the two brain regions, a continuous line represents significant (p < 0.05) correlations whereas a dashed line represents a trend to significant correlation. Two ensembles were identified in both groups. In blue, the regions vDG, dDG and dACC were present in both treatments, but the RSC and dCA3 were included only in the CNO all-days group. In brown, the vCA1 and vCA3 were present in both treatments, but the dCA3 was included in the Vehicle group and the BLA and dCA1 were included in the CNO all-days. (E) Heatmap matrix representing interregional results from the Fisher z-transform test for differences between two correlations. Z-scores values range from +2 to −4. Positive z-scores refer to higher rho coefficient value in the Vehicle group compared to CNO all-days. Negative values refer to higher coefficient in the CNO all-days compared to Vehicle. Values above +1.96 or below −1.96 are considered significant. CNO all-days has higher correlations between the dCA3-RSC, dCA2-vDG and BLA-vCA1 compared to Vehicle. (F) Central hubs for the Vehicle (left) and CNO all days (right) groups were determined as the intersection between the top 10% of regions with highest scores of degrees (pink), betweenness (blue) and eigenvectors (green) The dDG and vCA1 were considered central hubs for the Vehicle group but the BLA was the only one in the CNO all-days group. All data are presented as mean ± SEM. n.s. = non-significant, *p < 0.05 when performing Spearman pairwise correlations, #p<0.05 when comparing between treatment. (dDG) dorsal Dentate Gyrus, (dCA3) dorsal CA3, (dCA2) dorsal CA2, (dCA1) dorsal CA1, (BLA) basolateral nucleus of the Amygdala, (LS) lateral Septum, (MS) medial Septum, (RSC) Retrosplenial Cortex, (ACC) anterior Cingulate Cortex, (vDG) ventral Dentate Gyrus, (vCA3) ventral CA3, and (vCA1) ventral CA1.

## DISCUSSION

In this study, we established a robust model of rescue behavior in mice and, for the first time, identified a stable dHPC neural ensemble linked to prosocial actions, which persists across diverse spatial, temporal, and social contexts. These findings highlight the dHPC as a key hub in coordinating the interaction between cognitive and emotional networks, a process essential for the expression of prosocial actions.

Our findings demonstrate that emotional arousal in distressed conspecifics enhances rescue behavior in mice, with observers in the cold water (CW) condition exhibiting significantly shorter liberation latencies and higher success rates compared to the social separation (SS) only group. This aligns with rodent studies where heightened negative valence in victims accelerates prosocial responses, postulating increased empathy on the helper’s side^17–19^. A recent study^64^ demonstrated that rats exhibiting rescue behavior generally display higher levels of affiliative behavior. Future research should explore which additional behavioral traits—such as cognitive abilities and emotional responses—are associated with successful rescue behavior. Anxiety and stress seem to modulate prosocial behavior in human and non-human animals^65^. Chronic stress is well known to promote aggressive behavior and negatively impact brain function and cognition^66^. However, recent evidence suggests that acute stress, especially moderate levels of it, may promote prosocial behavior as well^67^. Anxiolytic treatment before helping behavior test sessions increased latency to freeing the caged partner and shifted the free rat’s motivation towards chocolate^35^. In contrast, rats that showed the highest corticosterone response to the HBT, when exposed to the trapped conspecific prior to training sessions, exhibited the longest latencies during subsequent testing. The authors proposed an inverted U-shaped relationship between stress response and rescue behavior in the HBT, wherein both low and high levels of emotional arousal impairs performance, while whereas moderate levels facilitate optimal helping behavior^35^. Another study also reported that acute, mild footshocks promoted helping behavior in a similar task to the HBT, whereas high intensity footshocks impaired it – strengthening the hypothesis of an inverted U-curve interaction^68^. The behavioral and neural mechanisms through which increased stress of the victim influences prosocial behavior of the observer remain a compelling subject for future investigation. Although we did not observe significant sex differences in the overall occurrence of rescue behavior, males and females appeared to employ different strategies. Emerging evidence highlights the relevance of sex differences in affiliative and prosocial behaviors^69^, warranting further investigation in this context. Moreover, while most mice showed longer latencies to liberate stranger victims compared to familiar ones, in several cases we observed markedly shorter latencies for specific stranger victims. This suggests that subtle, individual-specific social dynamics between the potential helper and the victim—beyond mere familiarity—may influence rescue behavior.

Silencing the dHPC—but not the vHPC—impaired the acquisition of helping behavior, as evidenced by a reduction in door-interaction attempts, without eliminating the ability to interact with the door, and by increased liberation latencies. This dissociation suggests a possible dHPC role in integrating episodic representations of spatial context (e.g., door location, victim position and spatial cues) with the affective salience of the victim’s distress, a process critical for forming adaptive prosocial intentions^45,57^. vHPC silencing spared learning but altered post-liberation social investigation, in line with its previously shown role in affective empathy and social memory consolidation^53,56^. Functional connectivity changes induced by dHPC silencing further suggest that disrupting hippocampal activity alters how the brain connects sensory, motivational, and contextual information. The functional recruitment of dorsal CA1 neurons during learning further supports the hypothesis that hippocampal cell ensembles encode task-relevant spatial and social contingencies, enabling mice to flexibly update their behavioral strategies across sessions. Our results align with the recently proposed broader role of the dorsal hippocampus in action planning^61,70^, position^71^ as well as sex, identity, and affiliation of another conspecific^72^. Future studies should further explore how actions with similar or divergent valences influence dHPC network recruitment, while distinguishing which components of these networks are specific to certain tasks and which are generalizable across different contexts.

Interestingly, the hippocampus may play a distinct role in social emotions like empathy compared to basic emotions, likely due to its involvement in relational thinking and autobiographical memory retrieval. In humans, hippocampal damage disrupts perspective-taking and empathic concern^44^, while episodic simulation of helping scenarios engages the medial temporal lobe (MTL) subsystem to generate prosocial intentions^55^. Our work parallels these findings, showing that dHPC-dependent spatial and relational coding in mice facilitates the association between rescue behavior and the alleviation of a conspecific’s distress—akin to the MTL’s role in constructing vivid, scenario-based simulations in humans. Furthermore, the behavioral impairments caused by dHPC inactivation resemble the social cognition deficits observed in patients with hippocampal damage. The interplay between the dHPC, vPC and the amygdala can be further elucidated through studies on hippocampal–amygdala memory circuits involved in experience-dependent observational fear^53^. Critically, the dissociation shown in our results between dHPC (action planning) and vHPC (post-liberation social engagement) mirrors the MTL’s dual role in episodic memory and emotional valence processing, respectively^50^. This aligns with the hypothesis that social emotions like empathy require hippocampal integration of self-referential memories with online social computations^73^. It is worthy of note that hippocampal engagement during social emotion regulation may involve dynamic interactions with regions like the vmPFC, anterior insula, retrosplenial and cingulate cortex, which are central to emotional experience and perspective-taking^74,75^.

Emerging evidence in humans suggests the association of different neurocognitive networks with the affective and cognitive components of empathy, the Salience and the Default Mode networks respectively^76^. The Salience network (SN) especially was associated with high scores in empathy tasks^77^. Recently, several studies started mapping homologous brain regions to these networks in rodents, and they found evidence that the Anterior Cingulate Cortex (ACC), Prelimbic cortex (PrL), anterior Insular Cortex (aIC), basolateral Amygdala (BLA) are also functionally connected in both mice and rats^78,79^. These regions are involved with emotional contagion and helping behavior in rodents, too. In particular, it was demonstrated that the ACC is necessary for emotional contagion^23^, that the aIC is involved in the emotional arousal needed to motivate helping^80^ and that the inactivation of the BLA impairs pro-social behaviors^81^. The vHPC is anatomically connected to the BLA, PrL, and aIC^49^, and the BLA-vHPC projections are involved in social memory and social behavior ^82^ as well as anxiety regulation^83^. In our chemogenetics experiments, vehicle-treated mice exhibited strong correlations between dHPC and vHPC subregions (DG, CA3, CA1) and ACC, suggesting an efficient network for adaptive prosocial behavior. Chemogenetic inactivation of the dHPC disrupted the coactivation profile, notably enhancing correlations between the BLA and the dHPC, vHPC, and ACC. This alteration positioned the BLA—a region implicated in emotional processing and defensive responses^47^—as a central hub. Future studies employing high-resolution, multi-region brain activity measurements should investigate how synchronization across different brain regions dynamically changes during prosocial behavior and its absence.

In summary, our study reveals a previously unrecognized role of dorsal hippocampal networks in regulating prosocial behavior. Impaired prosocial behavior frequently co-occurs with conditions such as antisocial personality disorder, psychopathic traits, depression, and autism^84^. Advancing treatment strategies for these disorders necessitates a fundamental understanding of the neuronal mechanisms underlying prosocial behavior. Rodent models serve as a powerful tool for this purpose, enabling high-resolution monitoring of brain activity at the cellular level and facilitating the investigation of distinct cell types’ contributions within relevant neural circuits. Insights gained from our results can facilitate the development of novel genetic and environmental models of human disorders associated with empathy deficits, ultimately paving the way for targeted therapeutic interventions.

## ACKNOWLEDGEMENTS

We dedicate this manuscript to Mateus dos Santos Corrêa, in memoriam. This work was supported by the Alexander von Humboldt Foundation (fellowship to MSC), Leibniz Best Minds StartUp grant (J143/2022), and German Center for Mental Health (DZPG, project 01EE2305E). We thank Daniela Hill for her technical assistance, and all members of the Cognition & Emotion Lab for their valuable discussions.

## AUTHOR CONTRIBUTIONS

Conceptualization, S.M.; formal analysis, M.S.C, A.Ag., C.O., A.B., E.T., M.L., L.M. with inputs from S.M. and P.B.; investigation, M.S.C, A.Ag., A.B., E.T., M.L. with inputs from S.M., P.B. and A.A.; resources, S.M., A.A. and P.B.; visualization, M.S.C, A.Ag., A.B., E.T., M.L; writing – original draft, M.S.C. and S.M. with inputs from all authors; supervision, S.M.

## DECLARATION OF GENERATIVE AI AND AI-ASSISTED TECHNOLOGIES IN THE WRITING PROCESS

During the preparation of this work the authors used Notion© and Perplexity© in order to improve the readability and language of the manuscript. After using this tool/service, the authors reviewed and edited the content as needed and take full responsibility for the content of the published article.

## A SUPPLEMENTAL INFORMATION

Document S1. Figures S1–S7

Video S1. Helping behavior task, related to Figures 1 and 2.

Video S2. Calcium imaging recording in dCA1, related to Figure 3.

## STAR*METHODS

### KEY RESOURCES TABLE

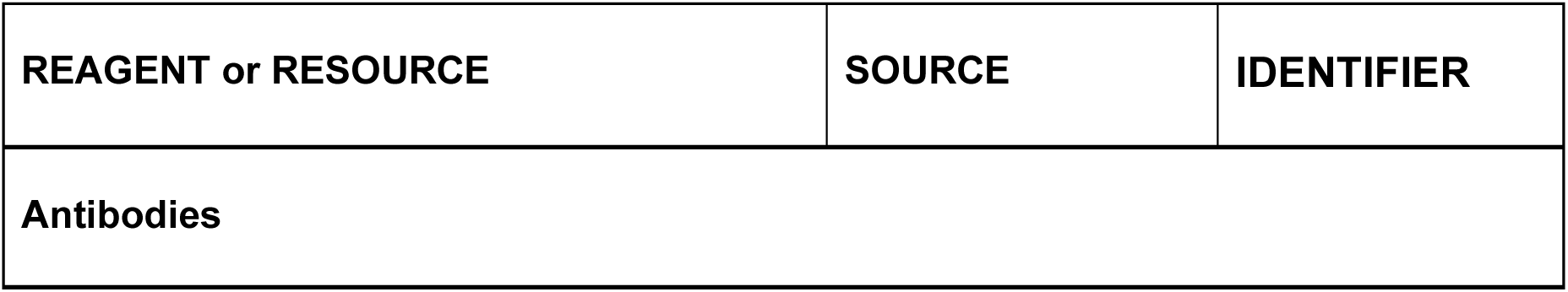
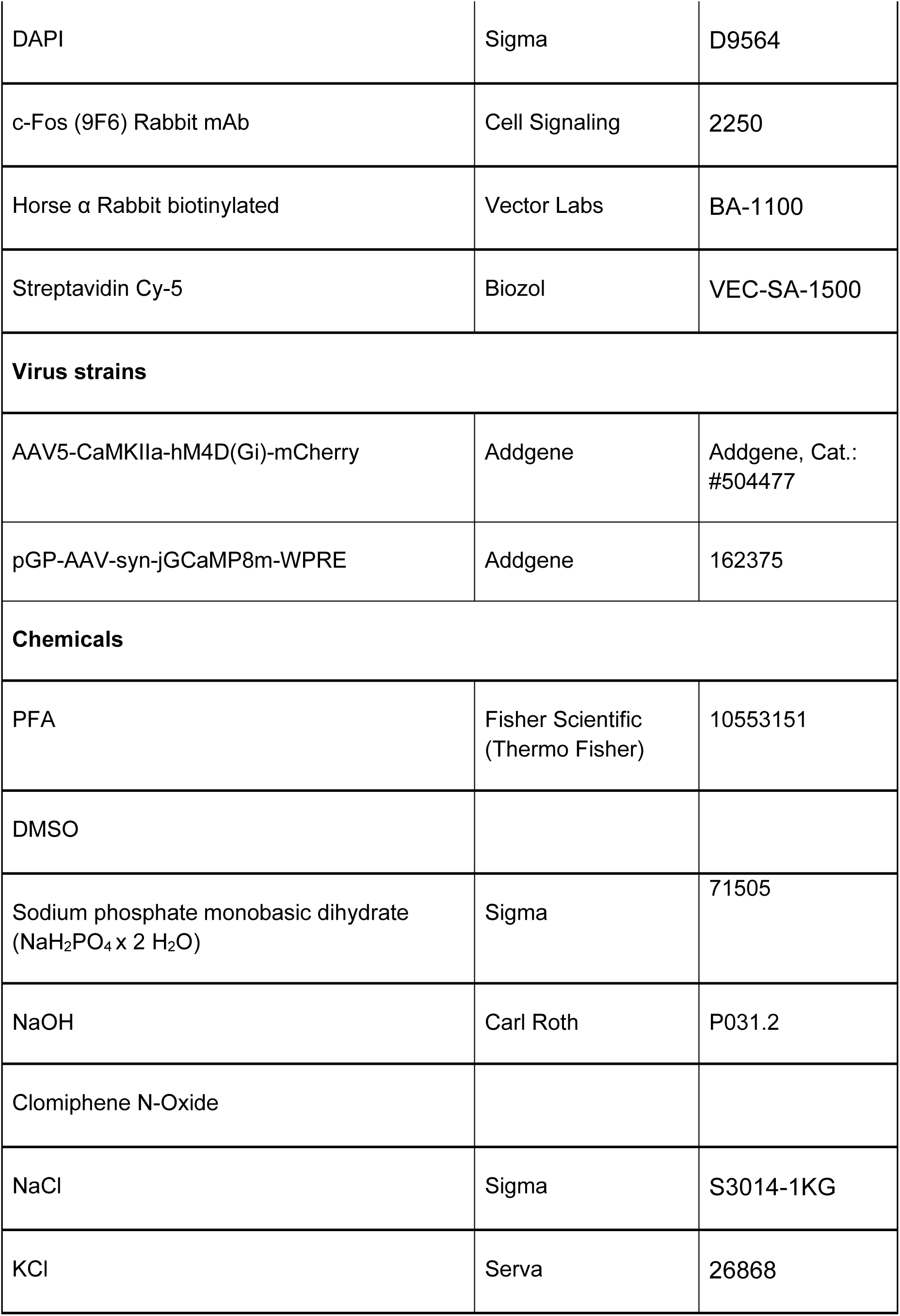
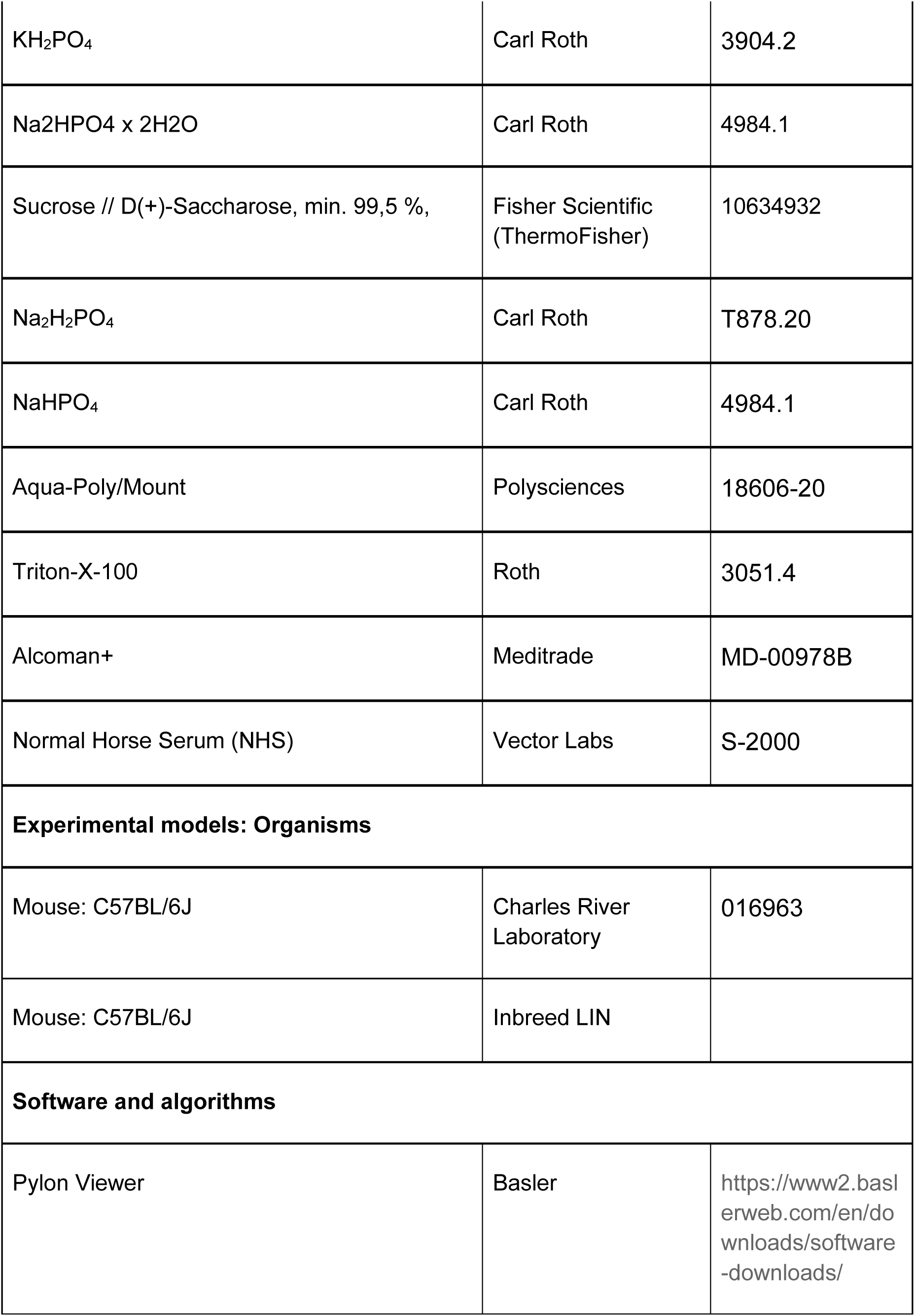
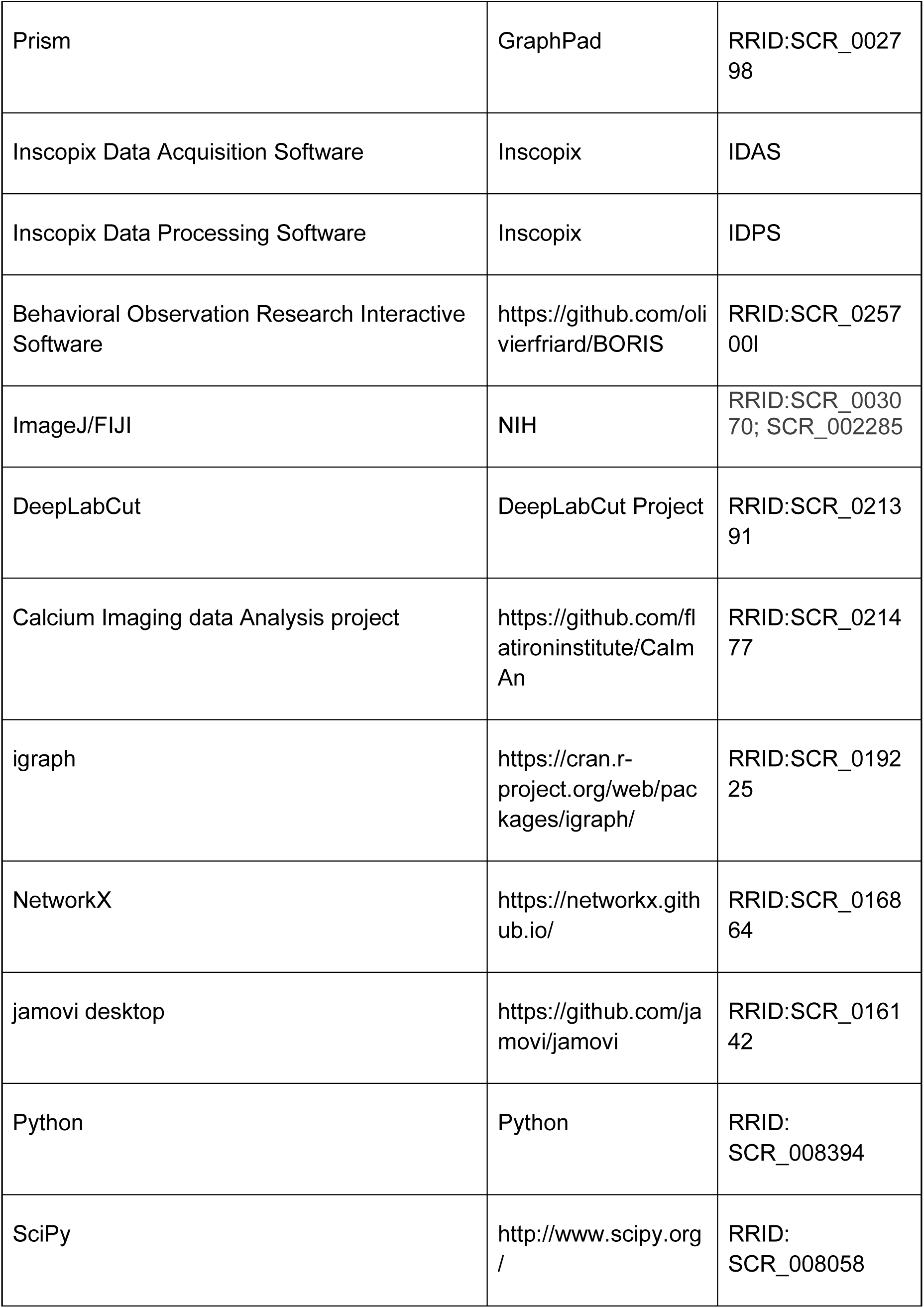
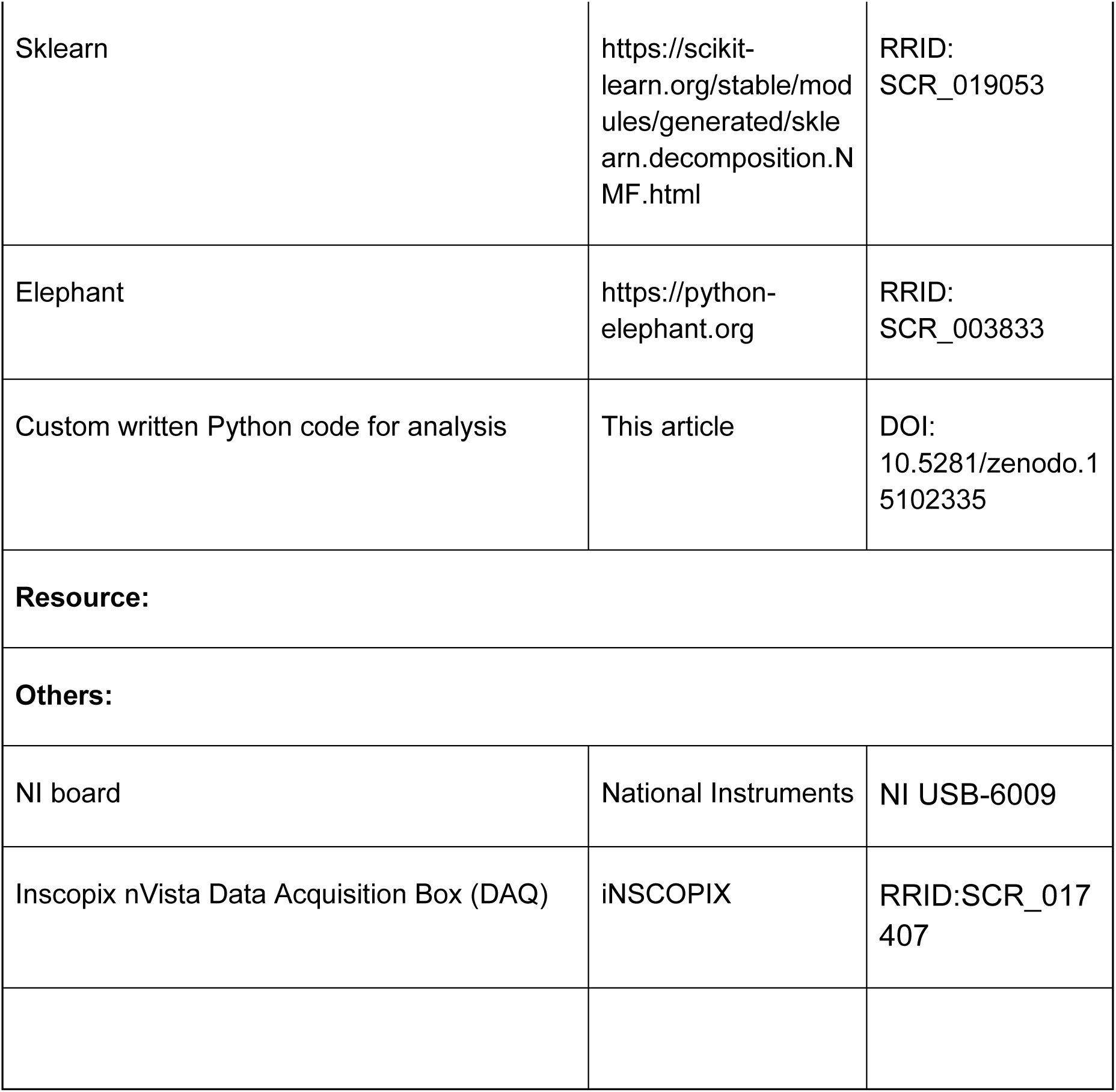

### CONTACT FOR REAGENT AND RESOURCE SHARING

Further information and requests for resources, reagents, and code should be addressed to and will be fulfilled by the Lead Contact, Sanja Mikulovic.

### EXPERIMENTAL MODEL AND SUBJECT DETAILS

Adult male and female mice (8 or 10 weeks old at the time of the surgery, or 12 weeks old for behavior only experiments) were maintained on a 12-h light/12-h dark cycle with ad libitum access to food and water. Subject mice used for all experiments were either from the C57BL/6J strain (Jackson Laboratories, 000664). Experiments were performed during the dark phase. All experiments were performed according the Directive of the European Communities Parliament and Council on the protection of animals used for scientific purposes (2010/63/EU) and were approved by the animal care committee of Sachsen-Anhalt, Germany. Subjects with mistargeted viral injections and/or GRIN lenses were excluded from analyses. In addition, miniscope mice with fewer than 50 accepted neurons on the first day of the training period were excluded from the analyses.

### METHOD DETAILS

#### Viral injections and surgical procedures

Mice were anesthetized with isoflurane and administered analgesics for all surgical procedures. A they were warmed with a 37°C heating pad (Fine Science Tools) throughout the procedure. The animals were head-fixed using a non-punctuated head bar and a nose clamp (MA-6N, Narishige), and placed under a motorized stereotactic frame (Luigs-Neumann). The eyes were kept moist with eye creme (Bepanthen). An incision was made in the scalp, and bregma and lambda were measured to ensure the skull was flat. A small craniotomy was performed on top of the region of interest and injections were administered using a 34-gauge needle and Hamilton syringe (World Precision Instruments). All coordinates were calculated using the Allen Mouse Brain Atlas as a reference. All viruses were injected using an UltraMicroPump (World Precision Instruments) at a rate of 150nL/min. After all injections the needle was left in place for at least 10 minutes before slow withdrawal. The scalp was then sutured, and the animals were injected i.p. with meloxicam (0.05 mg/kg) once daily for 3 days.

For the chemogenetic silencing of dHPC neurons, AAV-CaMKIIa-hM4D(Gi)-mCherry (Addgene, 504477-AAV5) was injected bilaterally into 10-week-old animals. A volume of 300 nl of virus (1.9 × 1012 pp ml−1) was injected per hemisphere at AP, −1.9 mm; ML, ±1.5 mm and DV, −1.4 mm from Bregma. For the chemogenetic silencing of vHPC neurons, the same procedure was performed, injecting a total volume of 500 nl of the same viral construct per hemisphere in two locations at AP, −3.2 mm; ML, ±3.5 mm and DV, −4.0 and −4.7 mm from Bregma.

For dCA1 miniscope implantation, pGP-AAV-syn-jGCaMP8m-WPRE (Addgene plasmid #162375-AAV1) expressing the genetically encoded calcium indicator GcaMP8m, was infused into the dCA1 region of 8 weeks old animals. A total volume of 500 nl (2.2e12 GC/mL) was injected unilaterally into the right hemisphere in two locations at AP, −1.9 mm; ML, +1.35 and +1.45 mm; DV, −1.3 from Bregma. Immediately after virus infusions, four points of 0.75 mm A, P, L, M from the given coordinates were marked. With the help of the marked points a circular craniotomy of ∼ 1.5 mm diameter was drilled. The piece of skull and the underlying dura was removed. The cortical tissue, along with horizontal and diagonal fibers of the corpus callosum were removed by using the aspiration technique, while cortex buffer was continuously flushed. A ProView integrated GRIN lens (diameter 1 mm, length 4.0mm, iNSCOPIX) was centered in the middle of the craniotomy and carefully moved down to DV: −1.40 (Figure 3 B). Finally, the lens was secured in place using dental cement (Gradia Direct Flo BW) applied on the skull.

Once the experiments were concluded all mice were transcardially perfused, first with PBS, followed by 4 % PFA in PBS. The brains were extracted and post-fixed in 4 % PFA in PBS for at least 24 hours. Afterwards, the brain tissue was sliced, and the viral expression was confirmed under a confocal microscope.

#### Behavioral assays

Before any behavioral test, mice were handled by the experimenter for a week and habituated to the experimental room for an additional week. Prior to the start of habituation, randomly assigned observer mice were earmarked in the right ear for easy role identification.

##### Setup

Our HBT setup was adjusted from the setup from^19^. This setup consisted two separated, rectangular, plexiglass compartments (15 cm x 10 cm x 25 cm). One of the compartments was called the observer mouse’s box, and the other was the victim mouse’s box. Both boxes were covered with plexiglass lids and had a 10 cm large hole, perfectly lined up with each-other, where a swinging door was attached on the observer’s side. A 1 cm gap between the compartments allowed passage between them. Additionally, the observer ‘s box had a base elevation of 5 cm. This setup was placed on a moveable high stand with a transparent top. For video recording of the experimental procedure, two cameras (Basler acA2040-90umNIR) were used. One was installed at the bottom of the setup for tracking mice’s trajectories and position, and the second on the side for facilitating the identification of their behaviors. Camera’s position was fixed throughout the experiment. Furthermore, two infrared lamps (THORLabs LIU 850A and LIU 780A) were mounted underneath the plexiglass stand, using them as a light source for the cameras. Raw behavior video data were acquired and analyzed at a 60 Hz sampling rate using Pylon Viewer 64-bit (Basler v. 7.4). Recordings started simultaneously by synchronizing them via a NI board that received a trigger command from the Measurement & Automation Explorer software (National Instruments v. 20).

##### Helping Behavior Task without calcium imaging recordings

###### Habituation

Mice were placed in pairs at the observer’s box once a day to explore it freely for 10 min for three consecutive days unless specified differently. Here, the observer’s box was detached from the victim’s box, and the connection hole to the victim’s box door was completely covered. The box was cleaned with 70% ethanol between sessions and with a few minutes dry out before reuse. After habituation, all pairs were returned to the holding room.

###### Training

In order to create an aversive environment for the victim the victim’s box was filled for each session with fresh, cold water (∼ 24 °C) to a depth of 5 cm, matching the observer’s box base elevation (See Fig. S1 B, Video S1). For the “neutral” condition, no water was added to the victim’s box (See Fig. S1 A). Lights remained turned off for the whole session duration and mice were exposed only to ambient noise during training. Next, the camera recordings were started, and both naïve mice were placed in their respective boxes, always first the observer and then the victim order. Afterward, boxes were covered with their lid. Each session lasted a maximum of 17 minutes. 15 minutes were the maximum period for the observer to open the door and at least 1,5 minute was the reunion phase at observer’s box. For sessions in which the observer mouse did not open the door or did not sustain door opening behavior long enough for the victim to cross, the experimenter terminated the session by opening the door after 15 minutes, allowing the victim mouse to cross to the observer’s box. These sessions were called “no liberation” sessions. Sessions in which the observer successfully opened the door and allowed the victim to cross to the safe side were called “liberation” sessions. Latency to liberation was measured as the time from victim insertion into the victim’s box until the moment that the victim crossed its head across the door opening (See Fig. S1 C), with no liberation sessions receiving a 15 min latency. The observer mice that learned to open the door and allowed victim liberation for three consecutive days, distributed towards the end of the training period, were classified as “helpers” in the analysis. Whereas, the observer mice that opened the door randomly only on one or two specific days or never opened at all were called “non-helpers”. At the end of each session, recording was stopped and both mice were returned to their home cage. Both compartments were cleaned with 70% ethanol before and after each session. All pairs were carried back to the holding room at the end of the experiment day.

###### Stranger Test

After six days of HBT training, observer mice were tested with an unfamiliar, same-sex victim that had been previously trained in a different pair. The rest of the behavioral session was performed as described previously. Latency to liberation was measured as previously described and the reunion interval with the stranger victim lasted at least one minute. After testing, each mouse was returned to its original home cage. All pairs were carried back to the holding room at the end of the test day.

##### Helping Behavior Task with pharmacogenetic silencing of dHPC or vHPC

###### Habituation, training and drug injection

To silence the targeted regions, 3–4 weeks after stereotaxic viral injections of AAV5-CaMKIIa-hM4D(Gi)-mCherry, mice were habituated to scruffing and syringe handling for three consecutive days. During training days, naïve mice were injected 30 min before behavioral experiments intraperitoneally with 5 mg kg^−1^ with a volume of 10 mg/mL of the iDREADD agonist CNO (Cayman Chemical, 34233-69-7) and returned to their homecage until behavioral testing. A control group of animals received treatment with a vehicle solution (1% DMSO in saline). Mice from a third group received vehicle injections from days one to five. On day six (and also Stranger test day for vHPC animals) they were treated with CNO. The treatments were administered daily for 6 days.

###### c-Fos immunohistochemistry

60 min after the final behavioral test, mice were transcardially perfused with phosphate buffered 4% paraformaldehyde solution (PFA). Brains were removed and post-fixed in PFA overnight, then in wt/vol 20% sucrose at 4°C for freeze protection. Brains were then snap-frozen in methyl butane cooled by liquid nitrogen and cut into 30 μm thick, serial coronal sections at the level of the PFC, nucleus accumbens, posterior ACC/medial, lateral septum, retrosplenial cortex/ dorsal hippocampus/ amygdala and ventral hippocampus on a cryostat. Free-floating sections were blocked with 5% normal horse serum (NHS) and 0.3% triton in phosphate buffer (PB) for 1 h. Afterwards the slices were incubated with a rabbit antibody against c-Fos (1:1000, Cell signaling #2250, Danvers, MA, USA) in 5%NHS and 0.3% triton for 48 h. It followed an incubation with biotinylated horse anti-rabbit secondary antibody (1:200, Biozol Diagnostica Vertrieb, Eching, Germany) in PB with 0.2% Triton for 2h and streptavidin-Cy5 labeling (1:1000, Biozol Diagnostica Vertrieb, Eching, Germany) in PB for 30 min. Nuclei were visualized with DAPI (300 nM, Thermofischer, Waltham, MA, USA). Immunostained sections were imaged using a DMI6000 epifluorescence microscope (Leica, Wetzlar, Germany) in both hemispheres from 2-3 slices per animal and region. c-Fos-positive cells were counted manually within the selected brain areas and cell density was normalized to area size measured with ImageJ software. The cell counts per slice and hemisphere were averaged for each individual animal in each target area and used for statistical comparison between groups.

##### Helping Behavior Task with calcium imaging recordings

###### Setup

The previously described setup was slightly modified for calcium imaging (See Fig. S1, video S2). The cameras were connected to a computer and the iNSCOPIX nVista data acquisition box (DAQ box) (iNSCOPIX, CA), which got triggered simultaneously at the start of each session by the DAQ box. The CA2+ video recording was controlled via DAQ box using Inscopix Data Acquisition Software (v. 2.4 iNSCOPIX, CA). The videos were recorded with either 20 or 60 frames per second. A second computer was connected to the DAQ box through Wi-Fi and used to control the Miniscope settings and recordings. The second computer was an additional light source in the experimental room. Importantly, miniscope observer mice were initially housed together with their future victim pairs, but after lens implantation they were single-housed until the end of the experiment to prevent any damages on the implants.

###### Handling

Before the experiment started, the animals were handled by the experimenter. For one and a half week the handling followed the previously described procedure. After that, observer mice were habituated to the holding, restraining and attachment of the Miniscope procedure in the experimental room. For three days, the experimenter restrained the observers with one hand in the experimental room, held it for a few seconds and released it by opening the hand. The procedure was repeated five times before the animal was released back to its home cage.

###### Habituation

For the habituation period, the animals were transferred from the holding room to the experimental room, where they stayed in their homecages for one hour in the dark to habituate. Following this, the re-habituation of animals to each other, to the setup and of the observer to the miniscope started. It was performed once per day for five days. First, the observer was removed from the home cage, restrained in the hand, the protection cap was taken off and the miniscope dummy was attached. The observer and victim were placed into the helper’s compartment. The box was covered with the Plexiglas lid, and the animals explored the box and each other freely for 10 minutes. The victim was returned to the home cage. The dummy miniscope was detached from the observer and the animal was released back into the home cage. The box was cleaned with 70 % ethanol. After each day of habituation, the cages were brought back to the holding room.

###### Training

Before each experiment, the naïve animals were transferred to the experimental room and stayed there for 30 to 60 min to habituate to the room. The miniscope was attached to the baseplate on the observer’s head and the mouse was placed back into the home cage for ten minutes to get accustomed to the wire of the miniscope, while the experimenter set up the miniscope and the recording. After the 10 minutes, the observer was placed into the observer’s box and the recording was started. Following this, the victim was transferred into the victim’s box, which got covered with the lid and the timer of the session was started. The sessions were conducted and latencies recorded See Figure S1 D). After each trial, the miniscope was detached from the observer and the animals were placed back into their individual home cages. At the end of each experimental day, animals were returned to the animal holding room. In total, six days of training were performed for the Miniscope experiment.

###### Recent Test days

For this cohort of animals, two so called recent test days were conducted. They occurred 24 and 48 hours after the end of the training period. The first test day was with the stranger victim, as previously described. The next day the experimental setup was changed to create a novel spatial context (Figure S1E), allowing for a spatial context discrimination test while maintaining the movable door as a strong cue to the previous training. The list of changed features are as follows: cleaning of the box with disinfectant solution (Alcoman+, Meditrade), addition of 4 colored, paper shapes to each wall of the observer’s box, addition of a pattern made of tape on the Plexiglas lid, covering of the sliding door face with tape to block visual contact between the victim and the observer, turning the experimental room white lights on during the session, and turning on of white noise at 75dB during the whole duration of the test trial. The novel context test day was called Recent CtxB. Latency to liberation of the victim was measured in both recent test days. Animals were returned to the animal holding room after the end of each test day.

###### Victimless Control days

To experimentally compare differential neural activity between training and test days and the influence of a present victim, three victimless control days after the recent test days were performed. In all control days, the observer was placed in the observer’s box with the Miniscope attached. On the first control day (Toy day, Figure S1F), the victim’s box was filled with water and a commercial mouse toy, which did not move, release any sound or spread any odor was placed inside. On the second control day (Pool day, Figure S1G), the victim’s box was filled with water only. For the third control day (Empty day, See Figure S1H), the victim’s box remained completely empty, with no toy or water. No latency was measured during control days.

###### Remote Test days

To test remote memory on the already trained observer mice, they were tested again 28 days after the end of the training period. The first remote test day was in the novel spatial context, called RemoteCtxB, and the box and experimental room was modified as previously described. The second remote test day, called RemoteHBT, was in the exact training box and procedures took place as reported above. The familiar victim was used for both test days. Latency for liberation was measured during both sessions. At the end of the experiment, brains were collected after perfusion for histological analysis of lens implantation and virus spread.

### QUANTIFICATION AND STATISTICAL ANALYSIS

#### Behavioral Analysis

##### Animal trajectory tracking with DeepLabCut

Mice position detection and tracking was performed using DeepLabCut^85^ for experiments involving chemogenetic silencing of the hippocampus. We trained a ResNet50 model for 500,000 epochs using a training set of 570 example frames extracted from 38 diverse recordings across various experimental sessions. The model was trained to detect the following animal body parts and experimental chamber features: nose, front paws (left and right), body center, hind paws (left and right), tail base, tail tip, and all four corners of the chamber. Details of the training parameters are provided in the configuration files accompanying the analysis code (file: config.yaml). The model was then applied to all of the data, which was cropped to contain only the observer’s chamber (file: scale_analysis_oversubfolders.py). In some videos, the box was inverted during the experiment. In such cases, all detected body parts and chamber corners were inverted back to ensure that locations are comparable between sessions in later steps (file: swap_coords_inhb.py).

To additionally ensure compatibility between sessions, we aligned all detected body parts to a reference chamber with fixed dimensions — a 10×15 arbitrary unit rectangle corresponding to the real chamber dimensions in centimeters—using the detected chamber corners as alignment points (file: align.py). To enhance tracking reliability, we used the center of mass of the aligned body parts as the animal location point. It was calculated by taking a likelihood-weighted average of all detected body parts, excluding the tail tip due to frequent detection errors (file: generate_combined_dataframe.py). This location point was then used to calculate various locomotion metrics (file: compute_all_vars.py). To achieve this, we first divided the data into running and resting bouts based on instantaneous speed statistics. A running bout was identified when the squared normalized velocity exceeded a specific threshold:

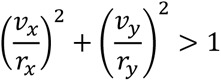

Here, 𝑣_𝑥_ and 𝑣_𝑦_ represent instantaneous velocities in X and Y directions, while 𝑟_𝑥_ and 𝑟_𝑦_ were calculated as a multiple 𝜆 = 5 of the median standard deviation of the instantaneous speed in X or Y directions. Running bouts thus determined the periods of active animal movement. The durations of these periods were summed and normalized by the length of the experimental session.

##### Manual behavioral annotation of videos

The latency to rescue was manually measured and defined as the time the observer took to liberate the victim (i.e., door opening by the observer and victim crossed its head across the door opening between the boxes). The timer started manually once the victim was inserted in the victim’s box and stopped when the victim crossed their head through the gap.

In addition, manual annotation of the chemogenetic silencing experiment was performed using the behavioral videos and the BORIS software. For BORIS, the main behaviors were specified with the help of modifiers and distinguished among them while watching the videos. The annotated behaviors were extracted from BORIS to a CSV file for each session and analyzed with custom written Python script (files: dHPC_plot.py, vHPC_plot.py). For each of the annotated behaviors, the mean behavior occurrence rate per minute was calculated as 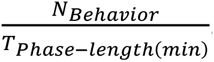, where 𝑁_𝐵𝑒ℎ𝑎𝑣𝑖𝑜𝑟_ is the total number of occurrences of a behavior and 𝑇_𝑃ℎ𝑎𝑠𝑒−𝑙𝑒𝑛𝑔𝑡ℎ(𝑚𝑖𝑛)_ is the total time in minutes before or after successful victim liberation. The annotated behaviors are described as follows:

1. Active door behavior before reunion (ADB): from the moment the victim is inserted in the victim’s box up to the moment that the victim crosses the head to the observer’s box, every instance in which the observer mouse interacted with the door using either its paws or snout was annotated as ADB.
2. Passive door behavior before reunion (PDB): during the same time interval from ADB, all the instances the observer spent exploring the door area by sniffing towards the door or directing its head to the door in close proximity to it (2 cm), without physically touching the door itself.
3. Door related behavior after reunion (DBA): after victim crossing, we annotated the time the observer mouse spent exploring the door area or gap between the boxes, either passively or actively.
4. Social behavior after reunion (SB): after victim crossing, we annotated the time the observer mouse spent exploring the victim (i.e., anogenital sniffing, snout sniffing, etc.).

#### Network generation and analysis for c-Fos immunohistochemistry

##### Network creation

We assessed neuronal activity and functional connectivity in response to HBT by examining changes in the expression of c-Fos, a product of the immediate early gene FOS and a marker of neuronal activation, in the chemogenetic silenced group of animals as previously described^42,70,86^. After eliminating extraneous entries, bilateral density values of c-Fos expression were obtained for each region of interest (ROI) by averaging hemispheric entries for individual animals. Within each experimental group (vehicle or CNO-all days), all possible pairwise correlations between c-Fos signals in the 12 analyzed brain regions were determined using Spearman correlation coefficients. These correlations, derived from vectors of sample size 5-9, were displayed as color-coded correlation matrices. Functional connectivity was defined when the Spearman’s rank correlation coefficient met the threshold of p ≤ 0.05 and rho ≥ 0.66. Then an identity matrix was built with values of 1 for each above-threshold correlation. To model c-Fos density data as a correlational network, we defined a graph structure G as the tuple of vertices and edges (V, E), where V represents brain regions, and E represents functionally connected pairs from the adjacency matrix A. This unweighted, undirected graph can be expressed mathematically as:

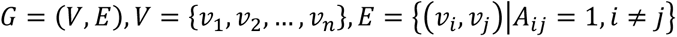

The effects of chemogenetic silencing of the dHPC on individual ROIs were investigated by contrasting the correlation of each treatment group using the Fisher r-to-z transformation, represented as a color-coded matrix. For hub identification, a series of centrality metrics for individual brain regions were examined^87^. This included degree (number of edges connected to a node), closeness (average length of the shortest possible path between a node and all others), eigenvector (frequency of a node’s connection to highly connected nodes), and clustering coefficient (frequency of a node’s connection to another node forming a local clique). Nodes falling in the top 10% of degree, closeness and eigenvector, as well as in the bottom 20% of the clustering coefficient, received a +1 “hub score” for each met criterion. Central hubs were defined as nodes with a hub score of 3 or greater (file: cFOS_HBT.ipynb).

#### Analysis of miniscope data

##### Preprocessing of one-photon calcium data

Raw calcium data were acquired at sampling rates of 60 or 40 Hz and converted to TIFF files using Inscopix Data Acquisition Software (v. 1.9 Inscopix). After conversion, the videos were downsampled to a uniform rate of 30 Hz in the temporal dimension and reduced by a factor of 2 in spatial dimensions to facilitate processing, using a custom Python script (Video S2, file: downsample.py). They were then cast to 16-bit format (file: cast_to_16.py). .. Cell detection was performed using CaImAn^88^ employing rigid motion-correction, CNMF-E model fitting and component evaluation (file: batch_cnmfe.py). The motion correction and CNMF-E model parameters are provided in the configuration files accompanying the analysis code (file: caiman_parameters.yaml). After manual inspection of the results, we selected parameters of *min_SNR*: 3.0, *SNR_lowest*: 2.5, *rval_thr*: 0.87 and *rval_lowest*: 0.4 for component evaluation with the aim to minimize false positive and negative components (file: reevaluate_comps.py). We removed duplicates (min_dist: 12) and all abnormally sized components (min_size_neuro: 5, max_size_neuro: 45). For the all subsequent analysis only accepted components were included, and for those the dF/F signals were detrended and z-scored (file: to_npy.py). The accepted components are referred to as “neurons” throughout the study. The percentage of neurons (% of neurons out of the total) was calculated as the number of neurons in a given session divided by the total number of neurons across all sessions.

##### Calcium Event Rate

The calcium event rate, defined as the number of calcium events per second, was determined using a peak detection algorithm applied to each neuron’s signal. To ensure robust detection, peaks were required to exceed a noise threshold and meet predefined criteria for event spacing (≥3 times the transient duration), width (equal to estimated transient duration), and height (≥1.5 times the estimated noise level). The transient duration was estimated from the data and the value of 400 ms was used. The event rate was then calculated as the number of detected peaks divided by the total signal duration in seconds (file: event_rate.py).

##### In silico data

To validate our pipeline and functional connectivity analysis results, we created an *in silico* dataset mimicking calcium imaging recordings of neuronal activity during prosocial behavior (file: generator.py). The parameters used to generate the in silico dataset are provided in the file accompanying the generating script (file: config.yaml). The simulated dataset was created to closely match the real experimental design, spanning a 6-day experimental period. The in silico dataset excluded control days and mice were categorized into helpers (80%) and non-helpers (20%). While non-helpers never initiated door-opening behavior, helpers showed probabilistic but always decreasing door-opening responses. On the first day, the helper’s door-opening latency followed a normal distribution with a mean maximum of 990 seconds. This latency progressively decreased across experimental days, as the mean was scaled by the experimental day (See Fig. S2 B).

Neurons were generated within a fixed 640 × 400-pixel frame to match the real data dimensions. The number of neurons ranged from 100 to 1000, appearing as a blob-shaped region with randomly assigned (X, Y) coordinates on the first experimental day, forming the spatial matrix A. The neuron body followed a Gaussian distribution centered in the field of view, with neuron diameters ranging from 8–12 pixels. Between sessions, neurons showed minor positional shifts limited to ≤10 pixels in any direction, to simulate natural variations.

For each neuron, we generated an individual rate profile containing random events of increased firing rates. We calculated the number of events by multiplying the session duration by the event rate sampled from normal distribution with the mean: 0.04–0.09 events per second and standard deviation: 0.001 or 0.01. We then generated event trains using an inhomogeneous Poisson process based on this rate profile, and convolved these events with an exponential decay function of the form:

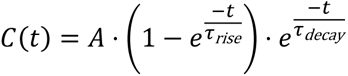

where 𝐴 (amplitude) ranged from 0.8 to 1.2, 𝜏_𝑟𝑖𝑠𝑒_ from 0.2 to 0.5, and 𝜏_𝑑𝑒𝑐𝑎𝑦_ from 1.5 to 3. These parameters were carefully chosen to match the temporal dynamics of real calcium transients. This process created calcium transients that formed the temporal matrix T. Since we generated event trains independently, any correlations between the resulting signals are purely chance occurrences.

We simulated imaging artifacts by adding light distortion effects (with intensity peaking at the center of the field of view) and Gaussian noise. The recordings were saved at 30 fps, with sessions lasting either 120 seconds after door opening or fixed 990 seconds (16.5 minutes) if the door remained closed. We generated the final calcium imaging data by multiplying each neuron’s spatial matrix (A) with its temporal matrix (T), storing the results as 3D TIFF image stacks. These in silico recordings were processed using the same pipeline as our real data to validate our observations and verify the analysis pipeline’s reliability (see subsections “Preprocessing of one-photon calcium data” and “Calcium Event Rate”).

##### Functional Connectivity Analysis

For functional connectivity analysis, calcium traces for each detected cell in the real and in silico datasets were extracted and preprocessed as specified in the previous section (file: functional_connectivity_M.py). To mitigate for any session length effect, pairwise correlations between cell activity signals by segmenting the ΔF/F signal in 1-minute interval bins and computing the Pearson correlation coefficients for each bin. The end result was calculated averaging the coefficients across the whole session. These averaged coefficients resulted in a pairwise correlation matrix. We set a threshold of 0.3 to further create an adjacency matrix with values of 1 to each above threshold correlation. The threshold was selected manually to avoid potential ceiling effects and was compared to the in silico data. This approach ensured the inclusion of neuronal pairs with Pearson Correlation Coefficients above chance level (See Fig. S4 C). This allowed us to construct functional connectivity matrices representing the network structure of the imaged neuronal population where the connection between different nodes corresponds to the correlation value > 0.3 between given neurons. Further analysis of the functional connectivity data included measurement of average degree per node, clustering coefficient and global efficiency of the given network that was done using *networkx* library in Python.

For community analysis, all positive and negative Pearson correlation coefficients became the edges between nodes, meaning that the correlation coefficients were used as the weights associated between nodes. Intra-network communities (also referred to as ensembles) were identified by the Leiden algorithm ^89^. For our implementation, we used the *leiden_communities()* method in *igraph* (ver. 10.0.4) using CPM as the objective function. We did a parameter search over 10,000 values between 0.5 and 1.75 to find the ideal resolution parameter. Whichever parameter yielded the highest modularity was selected, after which the Leiden algorithm was run again for 10,000 iterations for all graphs in our analysis. Number of ensembles that were formed by 2 or more neurons per mouse and the average size of ensembles, also per mouse, excluding all ensembles composed by a single neuron were extracted as community related metrics.

For community Multi-Scale Sample Entropy (MSE), we employed a coarse-graining method across multiple scales ranging 1-9 ^90^. This method evaluates the complexity of a time series by quantifying its entropy over a range of temporal scales. This involved averaging the signal over different time scales to capture its temporal dynamics. The entropy values for each scale were calculated using a standard entropy algorithm (*SampEn*), allowing us to assess the complexity of neuronal activity. A value of 0 represents a regular and periodic signal whereas uncorrelated random signals will show higher values. We calculated entropy values using scales=9, m=2, and r=0.15, to quantify the complexity of the ensemble averaged signal during the whole session length.

##### Identification of liberation ensembles and synchronization metrics

Following the identification of neuronal ensembles, we examined their responses to liberation events, assessing calcium activity around the liberation time (file: liberation_communities_analyzer.py). For each ensemble, we extracted the averaged calcium signal for all neurons in a window size of 34 seconds (17s before and 17s after the victims head crossing) and compared the ensemble signal to the rest of the session. The window size of 17s was determined based on visual inspection of raw data. We proceeded to extract metrics comparing the event window with the rest of the signal: 1) ensemble MSE as previously described, 2) liberation calcium event AUC (Area Under the Curve), 3) neuronal synchronization (the averaged Pearson coefficient of all pairwise correlations between the calcium traces of neurons in the ensemble), and 4) ratio between neuronal synchronization during the liberation interval compared to the synchronization in the rest of the signal (positive values indicate higher synchronization of neuronal ensemble during liberation compared to the rest of the session). After extracting general features of the ensemble signals, we investigated whether some ensembles respond more strongly and specifically to the liberation event and whether these so called ‘liberation ensembles’ show differential features compared to the ‘non-liberation’ ones.

To do this, we calculated the likelihood that the observed liberation event-related activity occurred by chance. We normalized the ensemble averaged calcium signal and isolated segments into two groups, depending if they belong to the liberation window or the remaining signal. We proceeded to calculate the AUC above noise level for each peak during the liberation window and during the rest of the session. To account for uneven session durations, the background signal was segmented into non-overlapping windows matching the liberation event window. Finally, the event specificity was quantified using a Poisson test, comparing the observed event rate to the expected rate based on the mean background activity rate (calculated using a window-based background rate). Ensembles were classified as ‘liberation ensembles’ if they showed event-specific responses exhibiting a significantly higher activity rate during liberation event (p-value ≤ 0.00001). After identifying the ‘liberation’ ensembles, we compared the previously described signal features between the liberation and non-liberation ensemble sets to assess differences in their functional roles.

For peak detection, a fixed threshold was calculated using noise statistics (median of all signal values up to 65oth percentile + 2 * standard deviation). Peaks were detected in both event and background signals when considered higher than noise level. Criteria were the following: minimum length of peak = 200ms, maximum length of peak = 5s, height = noise level, prominence = 0.35 and minimum distance between peaks = 125ms.

#### Statistical analysis

Statistical analyses were conducted using Prism 8 and JAMOVI. A general linear mixed model (GLMM) approach was applied to most datasets due to its suitability for handling both fixed and random effects, which aligns well with our repeated measures design and hierarchical data structure. The GLMMs were implemented using the GAMLj module in JAMOVI. Various factor coding schemes were employed to address specific research questions and enable interpretable contrasts across repeated measures and experimental conditions, with individual mice modeled as a random factor. To analyze sessions or days of helping behavior, we used difference coding (inverse Helmert coding), which compares each level to the mean of all previous levels and is particularly useful for assessing cumulative effects over time. Simple factor coding was used for comparisons between two categorical groups, such as experimental versus control conditions, allowing for direct contrasts between targeted conditions. For comparisons involving three categorical groups, we applied repeated factor coding to facilitate pairwise comparisons between each group and a designated reference group. In certain cases, Helmert coding was implemented to compare each condition to the average of subsequent levels, thereby capturing progressive changes across conditions.

Additional statistical tests were applied as appropriate: chi-square tests for categorical data, repeated measures one- or two-way ANOVAs followed by Holm-Sidak post-hoc analysis, and two-sample Kolmogorov-Smirnov tests with bootstrapping (n = 100 iterations, resampling 150 data points per group without replacement). Where applicable, multiple comparisons were corrected using the Bonferroni method. For non-parametric two-sample comparisons, we used the Mann-Whitney test, and the Kruskal-Wallis test followed by Dunn’s post-hoc test was used for multiple group comparisons. Pearson correlations were calculated for continuous variables to assess linear relationships. All data are reported as mean ± SEM unless otherwise specified. Statistical significance was determined at a threshold of p < 0.05.

#### Python packages

The majority of the code was implemented in Python and makes use of the following packages: ‘os’ (v2.1.4), ‘sys’ (v3.11.4) and ‘utils’ (v1.0.2) for system administration; ‘numpy’ (v2.0), ‘scipy’ (v1.14.0) and ‘pandas’ (v2.2.2) for data processing, data analysis and general data handling; ‘scikit-learn’ (v1.5.0) for preprocessing of the data for classification and k-means clustering; ‘TensorFlow’ (v2.11) and ‘libsvm’ (v3.23.0.4) for LSTM and SVM classification, respectively; ‘Matplotlib’ (v3.9), ‘seaborn’ (v0.11.2) and ‘plotly’ (v5.7) for data visualization; ‘networkx’ (ver. 2.8.4), ‘statsmodels’ (ver. 0.13.2), ‘sklearn’ (ver. 1.2.1), ‘markov_clustering’ (ver. 0.0.06) and ‘igraph’ (ver. 10.0.4) for Network analysis

## DATA AND SOFTWARE AVAILABILITY

The data and software are available upon request to the corresponding author.

## REFERENCES

1. Panksepp, J.B., and Lahvis, G.P. (2011). Rodent empathy and affective neuroscience. Neurosci. Biobehav. Rev. 35, 1864–1875. 10.1016/j.neubiorev.2011.05.013.

2. Panksepp, J., and Panksepp, J.B. (2013). Toward a cross-species understanding of empathy. Trends Neurosci. 36, 489–496. 10.1016/j.tins.2013.04.009.

3. Gachomba, M.J.M., Esteve-Agraz, J., and Márquez, C. (2024). Prosocial behaviors in rodents. Neurosci. Biobehav. Rev. 163. 10.1016/j.neubiorev.2024.105776.

4. Keysers, C., Knapska, E., Moita, M.A., and Gazzola, V. (2022). Emotional contagion and prosocial behavior in rodents. Trends Cogn. Sci. 26, 688–706. 10.1016/j.tics.2022.05.005.

5. Cuff, B.M.P., Brown, S.J., Taylor, L., and Howat, D.J. (2016). Empathy: A review of the concept. Emot. Rev. 8, 144–153. 10.1177/1754073914558466.

6. Nowbahari, E., Scohier, A., Durand, J.L., and Hollis, K.L. (2009). Ants, Cataglyphis cursor, use precisely directed rescue behavior to free entrapped relatives. PLoS One 4, 4–7. 10.1371/journal.pone.0006573.

7. Watanabe, S., and Ono, K. (1986). An experimental analysis of “empathic” response: Effects of pain reactions of pigeon upon other pigeon’s operant behavior. Behav. Processes 13, 269–277. 10.1016/0376-6357(86)90089-6.

8. Edgar, J.L., Lowe, J.C., Paul, E.S., and Nicol, C.J. (2011). Avian maternal response to chick distress. Proc. R. Soc. B Biol. Sci. 278, 3129–3134. 10.1098/rspb.2010.2701.

9. Lalot, M., Delfour, F., Mercera, B., and Bovet, D. (2021). Prosociality and reciprocity in bottlenose dolphins (Tursiops truncatus). Anim. Cogn. 24, 1075–1086. 10.1007/s10071-021-01499-z.

10. Masilkova, M., Ježek, M., Silovský, V., Faltusová, M., Rohla, J., Kušta, T., and Burda, H. (2021). Observation of rescue behaviour in wild boar (Sus scrofa). Sci. Rep. 11, 1–9. 10.1038/s41598-021-95682-4.

11. de Waal, F.B.M. (2012). The Antiquity of Empathy. Science (80-.). 336, 874–876. 10.1126/science.1220999.

12. De Waal, F.B.M. (2008). Putting the altruism back into altruism: The evolution of empathy. Annu. Rev. Psychol. 59, 279–300. 10.1146/annurev.psych.59.103006.093625.

13. De Waal, F.B.M., and Preston, S.D. (2017). Mammalian empathy: Behavioural manifestations and neural basis. Nat. Rev. Neurosci. 18, 498–509. 10.1038/nrn.2017.72.

14. Decety, J., Bartal, I.B.A., Uzefovsky, F., and Knafo-Noam, A. (2016). Empathy as a driver of prosocial behaviour: Highly conserved neurobehavioural mechanisms across species. Philos. Trans. R. Soc. B Biol. Sci. 371. 10.1098/rstb.2015.0077.

15. Thomas Greene, J. (1969). Altruistic behavior In the albino rat. Psychon. Sci. 14, 47–48.

16. Rice, G.E., and Gainer, P. (1962). “Altruism” in the albino rat. J. Comp. Physiol. Psychol. 55, 123–125. 10.1037/h0042276.

17. Bartal, I.B.A., Decety, J., and Mason, P. (2011). Empathy and pro-social behavior in rats. Science (80-.). 334, 1427–1430. 10.1126/science.1210789.

18. Kitano, K., Yamagishi, A., Horie, K., Nishimori, K., and Sato, N. (2022). Helping behavior in prairie voles: A model of empathy and the importance of oxytocin. iScience 25, 103991. 10.1016/j.isci.2022.103991.

19. Sato, N., Tan, L., Tate, K., and Okada, M. (2015). Rats demonstrate helping behavior toward a soaked conspecific. Anim. Cogn. 18, 1039–1047. 10.1007/s10071-015-0872-2.

20. Hernandez-Lallement, J., Van Wingerden, M., Marx, C., Srejic, M., and Kalenscher, T. (2015). Rats prefer mutual rewards in a prosocial choice task. Front. Neurosci. 9, 1–9. 10.3389/fnins.2014.00443.

21. Márquez, C., Rennie, S.M., Costa, D.F., and Moita, M.A. (2015). Prosocial Choice in Rats Depends on Food-Seeking Behavior Displayed by Recipients. Curr. Biol. 25, 1736–1745. 10.1016/j.cub.2015.05.018.

22. Scheggia, D., La Greca, F., Maltese, F., Chiacchierini, G., Italia, M., Molent, C., Bernardi, F., Coccia, G., Carrano, N., Zianni, E., et al. (2022). Reciprocal cortico-amygdala connections regulate prosocial and selfish choices in mice. Nat. Neurosci. 25, 1505–1518. 10.1038/s41593-022-01179-2.

23. Hernandez-Lallement, J., Attah, A.T., Soyman, E., Pinhal, C.M., Gazzola, V., and Keysers, C. (2020). Harm to Others Acts as a Negative Reinforcer in Rats. Curr. Biol. 30, 949–961.e7. 10.1016/j.cub.2020.01.017.

24. Hess, E.M., Venniro, M., and Gould, T.D. (2023). Relative to females, male rats are more willing to forego obtaining sucrose reward in order to prevent harm to their cage mate. Psychopharmacology (Berl). 10.1007/s00213-023-06435-2.

25. Song, D., Wang, C., Jin, Y., Deng, Y., Yan, Y., Wang, D., Zhu, Z., Ke, Z., Wang, Z., Wu, Y., et al. (2023). Mediodorsal thalamus-projecting anterior cingulate cortex neurons modulate helping behavior in mice. Curr. Biol. 33, 4330–4342.e5. 10.1016/j.cub.2023.08.070.

26. Sun, F., Wu, Y.E., and Hong, W. (2025). A neural basis for prosocial behavior toward unresponsive individuals. Science 387, eadq2679. 10.1126/science.adq2679.

27. Zhang, M., Wu, Y.E., Jiang, M., and Hong, W. (2024). Cortical regulation of helping behaviour towards others in pain. Nature 626, 136–144. 10.1038/s41586-023-06973-x.

28. Netser, S., Meyer, A., Magalnik, H., Zylbertal, A., de la Zerda, S.H., Briller, M., Bizer, A., Grinevich, V., and Wagner, S. (2020). Distinct dynamics of social motivation drive differential social behavior in laboratory rat and mouse strains. Nat. Commun. 11. 10.1038/s41467-020-19569-0.

29. Pozo, M., Milà-Guasch, M., Haddad-Tóvolli, R., Boudjadja, M.B., Chivite, I., Toledo, M., Gómez-Valadés, A.G., Eyre, E., Ramírez, S., Obri, A., et al. (2023). Negative energy balance hinders prosocial helping behavior. Proc. Natl. Acad. Sci. 120, 2017. 10.1073/pnas.2218142120.

30. Ueno, H., Suemitsu, S., Murakami, S., Kitamura, N., Wani, K., Matsumoto, Y., Okamoto, M., and Ishihara, T. (2019). Helping-Like Behaviour in Mice Towards Conspecifics Constrained Inside Tubes. Sci. Rep. 9, 1–11. 10.1038/s41598-019-42290-y.

31. Hollis, K.L., and Nowbahari, E. (2013). Toward a behavioral ecology of rescue behavior. Evol. Psychol. 11, 647–664. 10.1177/147470491301100311.

32. Silberberg, A., Allouch, C., Sandfort, S., Kearns, D., Karpel, H., and Slotnick, B. (2014). Desire for social contact, not empathy, may explain “rescue” behavior in rats. Anim. Cogn. 17, 609–618. 10.1007/s10071-013-0692-1.

33. Hachiga, Y., Schwartz, L.P., Silberberg, A., Kearns, D.N., Gomez, M., and Slotnick, B. (2018). Does a rat free a trapped rat due to empathy or for sociality? J. Exp. Anal. Behav. 110, 267–274. 10.1002/jeab.464.

34. Schwartz, L.P., Silberberg, A., Casey, A.H., Kearns, D.N., and Slotnick, B. (2017). Does a rat release a soaked conspecific due to empathy? Anim. Cogn. 20, 299–308. 10.1007/s10071-016-1052-8.

35. Bartal, I.B.A., Shan, H., Molasky, N.M.R., Murray, T.M., Williams, J.Z., Decety, J., and Mason, P. (2016). Anxiolytic treatment impairs helping behavior in rats. Front. Psychol. 7, 1–14. 10.3389/fpsyg.2016.00850.

36. Peng, S., Li, M., Yang, X., and Xie, W. (2025). The neural basis of affective empathy: What is known from rodents. Neuropharmacology 269, 110347. 10.1016/j.neuropharm.2025.110347.

37. Paradiso, E., Gazzola, V., and Keysers, C. (2021). Neural mechanisms necessary for empathy-related phenomena across species. Curr. Opin. Neurobiol. 68, 107–115. 10.1016/j.conb.2021.02.005.

38. Bernhardt, B.C., and Singer, T. (2012). The neural basis of empathy. Annu. Rev. Neurosci. 35, 1–23. 10.1146/annurev-neuro-062111-150536.

39. Cox, S.S., Brown, B.J., Wood, S.K., Brown, S.J., Kearns, A.M., and Reichel, C.M. (2024). Neuronal, affective, and sensory correlates of targeted helping behavior in male and female Sprague Dawley rats. Front. Behav. Neurosci. 18. 10.3389/fnbeh.2024.1384578.

40. Yamagishi, A., Lee, J., and Sato, N. (2020). Oxytocin in the anterior cingulate cortex is involved in helping behaviour. Behav. Brain Res. 393, 112790. 10.1016/j.bbr.2020.112790.

41. Cox, S.S., Kearns, A.M., Woods, S.K., Brown, B.J., Brown, S.J., and Reichel, C.M. (2022). The role of the anterior insular during targeted helping behavior in male rats. Sci. Rep. 12, 1–11. 10.1038/s41598-022-07365-3.

42. Ben-Ami Bartal, I., Breton, J.M., Sheng, H., Long, K.L.P.L., Chen, S., Halliday, A., Kenney, J.W., Wheeler, A.L., Frankland, P., Shilyansky, C., et al. (2021). Neural correlates of ingroup bias for prosociality in rats. Elife 10, 1–26. 10.7554/eLife.65582.

43. Rubin, R.D., Watson, P.D., Duff, M.C., and Cohen, N.J. (2014). The role of the hippocampus in flexible cognition and social behavior. Front. Hum. Neurosci. 8, 1–15. 10.3389/fnhum.2014.00742.

44. Beadle, J.N., Tranel, D., Cohen, N.J., and Duff, M.C. (2013). Empathy in hippocampal amnesia. Front. Psychol. 4, 1–12. 10.3389/fpsyg.2013.00069.

45. Gaesser, B., Hirschfeld-Kroen, J., Wasserman, E.A., Horn, M., and Young, L. (2019). A role for the medial temporal lobe subsystem in guiding prosociality: the effect of episodic processes on willingness to help others. Soc. Cogn. Affect. Neurosci. 14, 397–410. 10.1093/scan/nsz014.

46. Sawczak, C., McAndrews, M.P., Gaesser, B., and Moscovitch, M. (2019). Episodic simulation and empathy in older adults and patients with unilateral medial temporal lobe excisions. Neuropsychologia 135, 107243. 10.1016/j.neuropsychologia.2019.107243.

47. Vieira, J.B., and Olsson, A. (2022). Neural defensive circuits underlie helping under threat in humans. Elife 11, 1–30. 10.7554/eLife.78162.

48. Scoville, W.B., and Milner, B. (1957). Loss of recent memory after bilateral hippocampal lesions. J. Neuropsychiatry Clin. Neurosci. 20, 11–21. 10.1136/jnnp.20.1.11.

49. Fanselow, M.S., and Dong, H.-W. (2010). Are the dorsal and ventral hippocampus functionally distinct structures? Neuron 65, 7–19. 10.1016/j.neuron.2009.11.031.

50. Strange, B.A., Witter, M.P., Lein, E.S., and Moser, E.I. (2014). Functional organization of the hippocampal longitudinal axis. Nat. Rev. Neurosci. 15, 655–669. 10.1038/nrn3785.

51. O’Keefe, J., Dostrovsky, J., and J. O’Keefe, J.D. (1971). Short Communications The hippocampus as a spatial map. Preliminary evidence from unit activity in the freely-moving rat. Brain Res. 34, 171–175.

52. Turner, V.S., O’Sullivan, R.O., and Kheirbek, M.A. (2022). Linking external stimuli with internal drives: A role for the ventral hippocampus. Curr. Opin. Neurobiol. 76, 102590. 10.1016/j.conb.2022.102590.

53. Terranova, J.I., Yokose, J., Osanai, H., Marks, W.D., Yamamoto, J., Ogawa, S.K., and Kitamura, T. (2022). Hippocampal-amygdala memory circuits govern experience-dependent observational fear. Neuron 110, 1416–1431.e13. 10.1016/j.neuron.2022.01.019.

54. Ben-Ami Bartal, I., Rodgers, D.A., Bernardez Sarria, M.S., Decety, J., and Mason, P. (2014). Pro-social behavior in rats is modulated by social experience. Elife 3, 1–16. 10.7554/elife.01385.

55. Gaesser, B., and Schacter, D.L. (2014). Episodic simulation and episodic memory can increase intentions to help others. Proc. Natl. Acad. Sci. U. S. A. 111, 4415–4420. 10.1073/pnas.1402461111.

56. Peng, S., Yang, X., Meng, S., Liu, F., Lv, Y., Yang, H., Kong, Y., Xie, W., and Li, M. (2024). Dual circuits originating from the ventral hippocampus independently facilitate affective empathy. Cell Rep. 43. 10.1016/j.celrep.2024.114277.

57. Keinath, A.T., Mosser, C.A., and Brandon, M.P. (2022). The representation of context in mouse hippocampus is preserved despite neural drift. Nat. Commun. 13. 10.1038/s41467-022-30198-7.

58. Liu, C., Todorova, R., Tang, W., Oliva, A., and Fernandez-Ruiz, A. (2023). Associative and predictive hippocampal codes support memory-guided behaviors. Science (80-.). 382. 10.1126/science.adi8237.

59. Dimsdale-Zucker, H.R., Ritchey, M., Ekstrom, A.D., Yonelinas, A.P., and Ranganath, C. (2018). CA1 and CA3 differentially support spontaneous retrieval of episodic contexts within human hippocampal subfields. Nat. Commun. 9. 10.1038/s41467-017-02752-1.

60. Pastalkova, E., Itskov, V., Amarasingham, A., and Buzsáki, G. (2008). Internally Generated Cell Assembly Sequences in the Rat Hippocampus. Science (80-.). 321, 1322–1327. 10.1126/science.1159775.

61. Zutshi, I., Apostolelli, A., Yang, W., Zheng, Z.S., Dohi, T., Balzani, E., Williams, A.H., Savin, C., and Buzsáki, G. (2025). Hippocampal neuronal activity is aligned with action plans. Nature, 2024.09.05.611533. 10.1038/s41586-024-08397-7.

62. Lee, H.S., and Han, J.H. (2022). Activity Patterns of Individual Neurons and Ensembles Correlated with Retrieval of a Contextual Memory in the Dorsal CA1 of Mouse Hippocampus. J. Neurosci. 43, 113–124. 10.1523/JNEUROSCI.1407-22.2022.

63. Yang, Y., and Wang, J.Z. (2017). From structure to behavior in basolateral amygdala-hippocampus circuits. Front. Neural Circuits 11, 1–8. 10.3389/fncir.2017.00086.

64. R. Hazani, J.M. Breton, E. Trachtenberg, B. Kantor, A. Maman, E. Bigelman, S. Cole, A.W., and I. Ben-Ami Bartal (2024). Helping behavior is associated with increased affiliative behavior, activation of the prosocial brain network and elevated oxytocin receptor expression in the nucleus accumbens.

65. Melloni, M., Lopez, V., and Ibanez, A. (2014). Empathy and contextual social cognition. Cogn. Affect. Behav. Neurosci. 14, 407–425. 10.3758/s13415-013-0205-3.

66. Beery, A.K., and Kaufer, D. (2015). Stress, social behavior, and resilience: Insights from rodents. Neurobiol. Stress 1, 116–127. 10.1016/j.ynstr.2014.10.004.

67. Buchanan, T.W., and Preston, S.D. (2014). Stress leads to prosocial action in immediate need situations. Front. Behav. Neurosci. 8, 1–6. 10.3389/fnbeh.2014.00005.

68. Karakilic, A., Kizildag, S., Kandis, S., Guvendi, G., Koc, B., Camsari, G.B., Camsari, U.M., Ates, M., Arda, S.G., and Uysal, N. (2018). The effects of acute foot shock stress on empathy levels in rats. Behav. Brain Res. 349, 31–36. 10.1016/j.bbr.2018.04.043.

69. Mikulovic, S., and Lenschow, C. (2025). Neural control of sex differences in affiliative and prosocial behaviors. Neurosci. Biobehav. Rev. 171, 106039. 10.1016/j.neubiorev.2025.106039.

70. Albrecht, A., Müller, I., Weiglein, A., Pollali, E., Çalışkan, G., and Stork, O. (2022). Choosing memory retrieval strategies: A critical role for inhibition in the dentate gyrus. Neurobiol. Stress 20. 10.1016/j.ynstr.2022.100474.

71. Omer, D.B., Maimon, S.R., and Las, L. (2018). Social place-cells in the bat hippocampus. 224, 6–11.

72. Ray, S., Yona, I., Elami, N., Palgi, S., Latimer, K.W., Jacobsen, B., Witter, M.P., Las, L., and Ulanovsky, N. (2025). Hippocampal coding of identity, sex, hierarchy, and affiliation in a social group of wild fruit bats. Science (80-.). 387. 10.1126/science.adk9385.

73. Immordino-Yang, M.H. (2016). Emotion, Sociality, and the Brain’s Default Mode Network: Insights for Educational Practice and Policy. Policy Insights from Behav. Brain Sci. 3, 211–219. 10.1177/2372732216656869.

74. Singer, T., and Lamm, C. (2009). The social neuroscience of empathy. Ann. N. Y. Acad. Sci. 1156, 81–96. 10.1111/j.1749-6632.2009.04418.x.

75. Lamm, C., Decety, J., and Singer, T. (2011). Meta-analytic evidence for common and distinct neural networks associated with directly experienced pain and empathy for pain. Neuroimage 54, 2492–2502. 10.1016/j.neuroimage.2010.10.014.

76. Betti, V., and Aglioti, S.M. (2016). Dynamic construction of the neural networks underpinning empathy for pain. Neurosci. Biobehav. Rev. 63, 191–206. 10.1016/j.neubiorev.2016.02.009.

77. Ziaei, M., Oestreich, L., Reutens, D.C., and Ebner, N.C. (2021). Age-related differences in negative cognitive empathy but similarities in positive affective empathy. Brain Struct. Funct. 226, 1823–1840. 10.1007/s00429-021-02291-y.

78. Tsai, P.J., Keeley, R.J., Carmack, S.A., Vendruscolo, J.C.M., Lu, H., Gu, H., Vendruscolo, L.F., Koob, G.F., Lin, C.P., Stein, E.A., et al. (2020). Converging Structural and Functional Evidence for a Rat Salience Network. Biol. Psychiatry 88, 867–878. 10.1016/j.biopsych.2020.06.023.

79. Zerbi, V., Floriou-Servou, A., Markicevic, M., Vermeiren, Y., Sturman, O., Privitera, M., von Ziegler, L., Ferrari, K.D., Weber, B., De Deyn, P.P., et al. (2019). Rapid Reconfiguration of the Functional Connectome after Chemogenetic Locus Coeruleus Activation. Neuron 103, 702–718.e5. 10.1016/j.neuron.2019.05.034.

80. Tomek, S.E., Stegmann, G.M., Leyrer-Jackson, J.M., Piña, J., and Olive, M.F. (2020). Restoration of prosocial behavior in rats after heroin self-administration via chemogenetic activation of the anterior insular cortex. Soc. Neurosci. 15, 408–419. 10.1080/17470919.2020.1746394.

81. Hernandez-Lallement, J., van Wingerden, M., Schäble, S., and Kalenscher, T. (2016). Basolateral amygdala lesions abolish mutual reward preferences in rats. Neurobiol. Learn. Mem. 127, 1–9. 10.1016/j.nlm.2015.11.004.

82. Felix-Ortiz, A.C., and Tye, K.M. (2014). Amygdala inputs to the ventral hippocampus bidirectionally modulate social behavior. J. Neurosci. 34, 586–595. 10.1523/JNEUROSCI.4257-13.2014.

83. Felix-Ortiz, A.C., Beyeler, A., Seo, C., Leppla, C.A., Wildes, C.P., and Tye, K.M. (2013). BLA to vHPC inputs modulate anxiety-related behaviors. Neuron 79, 658–664. 10.1016/j.neuron.2013.06.016.

84. Decety, J., and Moriguchi, Y. (2007). The empathic brain and its dysfunction in psychiatric populations: Implications for intervention across different clinical conditions. Biopsychosoc. Med. 1, 1–21. 10.1186/1751-0759-1-22.

85. Mathis, A., Mamidanna, P., Cury, K.M., Abe, T., Murthy, V.N., Mathis, M.W., and Bethge, M. (2018). DeepLabCut: markerless pose estimation of user-defined body parts with deep learning. Nat. Neurosci. 21, 1281–1289. 10.1038/s41593-018-0209-y.

86. Dos Santos Corrêa, M., Grisanti, G.D.V., Franciscatto, I.A.F., Tarumoto, T.S.A., Tiba, P.A., Ferreira, T.L., and Fornari, R.V. (2022). Remote contextual fear retrieval engages activity from salience network regions in rats. Neurobiol. Stress 18, 100459. 10.1016/j.ynstr.2022.100459.

87. van den Heuvel, M.P., and Sporns, O. (2013). Network hubs in the human brain. Trends Cogn. Sci. 17, 683–696. 10.1016/j.tics.2013.09.012.

88. Giovannucci, A., Friedrich, J., Gunn, P., Kalfon, J., Brown, B.L., Koay, S.A., Taxidis, J., Najafi, F., Gauthier, J.L., Zhou, P., et al. (2019). Caiman an open source tool for scalable calcium imaging data analysis. Elife 8, 1–45. 10.7554/eLife.38173.

89. Traag, V.A., Waltman, L., and van Eck, N.J. (2019). From Louvain to Leiden: guaranteeing well-connected communities. Sci. Rep. 9, 1–12. 10.1038/s41598-019-41695-z.

90. Humeau-Heurtier, A. (2015). The multiscale entropy algorithm and its variants: A review. Entropy 17, 3110–3123. 10.3390/e17053110.

91. de Melo, M.B., Daldegan-Bueno, D., Menezes Oliveira, M.G., and de Souza, A.L. (2022). Beyond ANOVA and MANOVA for repeated measures: Advantages of generalized estimated equations and generalized linear mixed models and its use in neuroscience research. Eur. J. Neurosci. 56, 6089–6098. 10.1111/ejn.15858.

92. Galucci, M. (2019). GAMLj: General analyses for linear models. at [jamovi module].

