## Supplementary Figures for "Neuronal networks in the dorsal hippocampus causally regulate rescue behavior in mice"

**SUPPLEMENTARY INFORMATION**

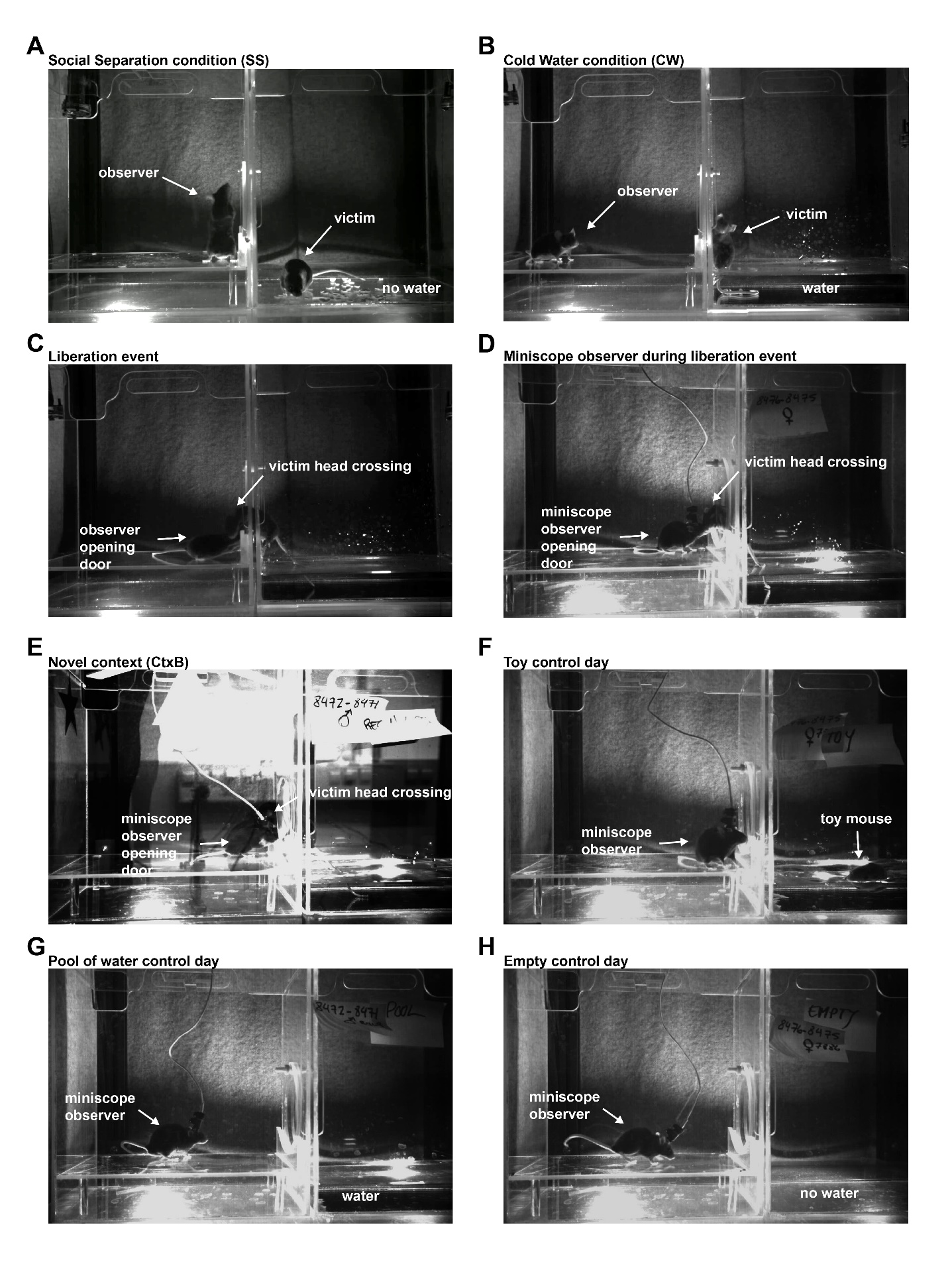

**Figure S1. Representative images illustrating different experimental conditions used in the study, related to Figure 1, 2, 3, 4.** (A) Social Separation (SS) condition: The setup consists of two interconnected chambers with a unidirectional door that only the observer mouse can open. A victim mouse is placed in the adjacent dry chamber. (B) Cold Water (CW) condition: Identical to (A), except the victim mouse is partially immersed in cold water in the adjacent chamber. (C) Liberation event: The moment of door opening by the observer mouse, allowing the victim to enter the observer’s chamber. The liberation event is precisely defined as the moment when the victim’s head fully crosses the chamber border. (D) Liberation event with miniscope recording: The observer mouse, equipped with a miniscope, opens the door while the victim crosses into the observer’s chamber. (E) Novel context (CtxB) condition: A liberation event occurs in a modified context, with the observer mouse equipped with a miniscope. The context was altered by cleaning with Alcoman+ disinfectant, adding colored paper shapes to the walls, placing patterned tape on the Plexiglas lid, covering the sliding door to block visual contact, turning on white room lights, and playing white noise (75 dB) during trials. (F) Toy control condition : Identical to (B), except a toy mouse is placed in the adjacent chamber and partially immersed in cold water instead of a live victim. (G) Pool of water control condition: Identical to (B), but without a victim mouse in the adjacent chamber containing cold water. (H) Empty control condition: Identical to (A), but with no victim mouse present.

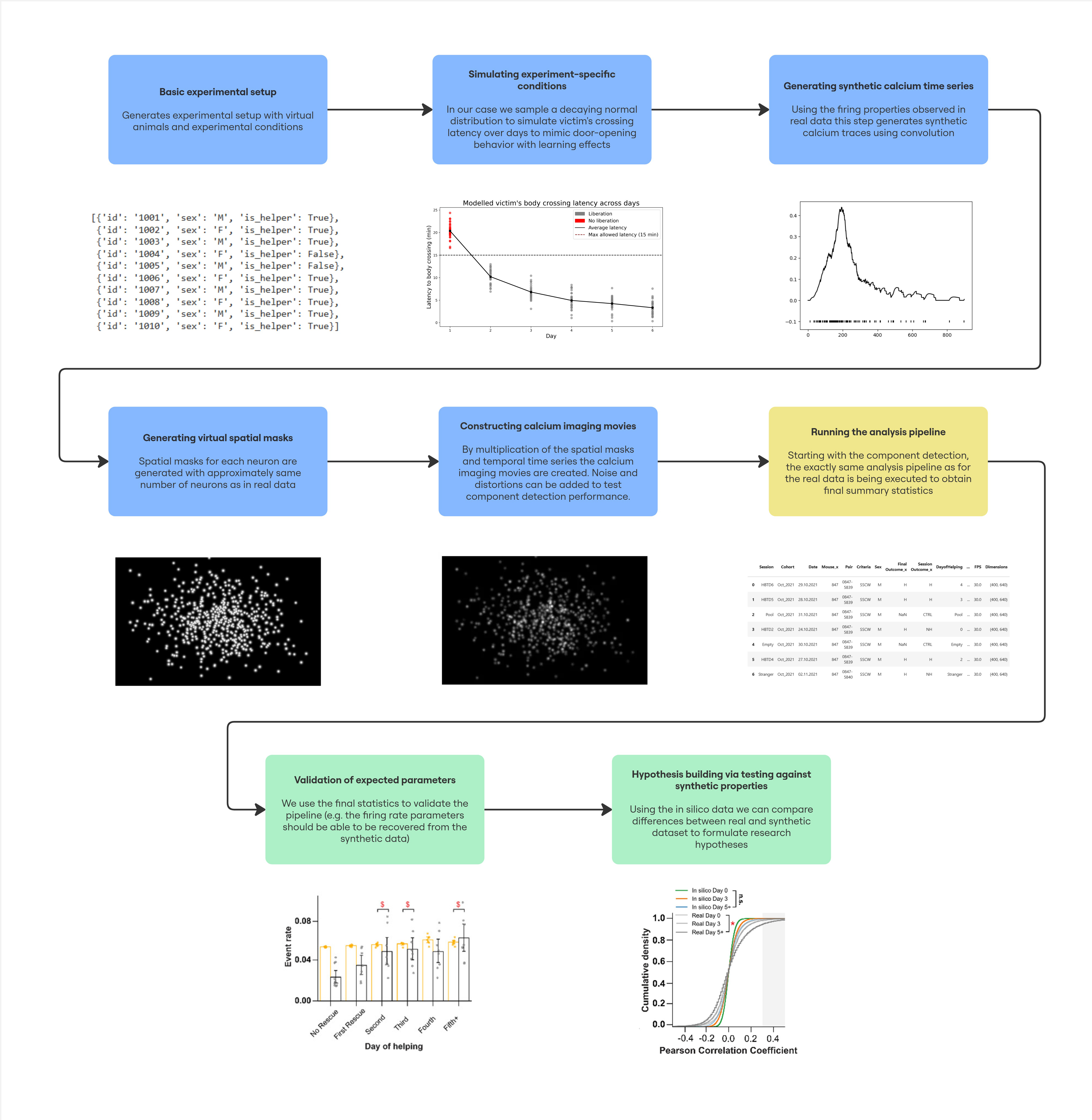

**Figure S2. Workflow for validating Calcium Imaging pipelines with synthetic data.**

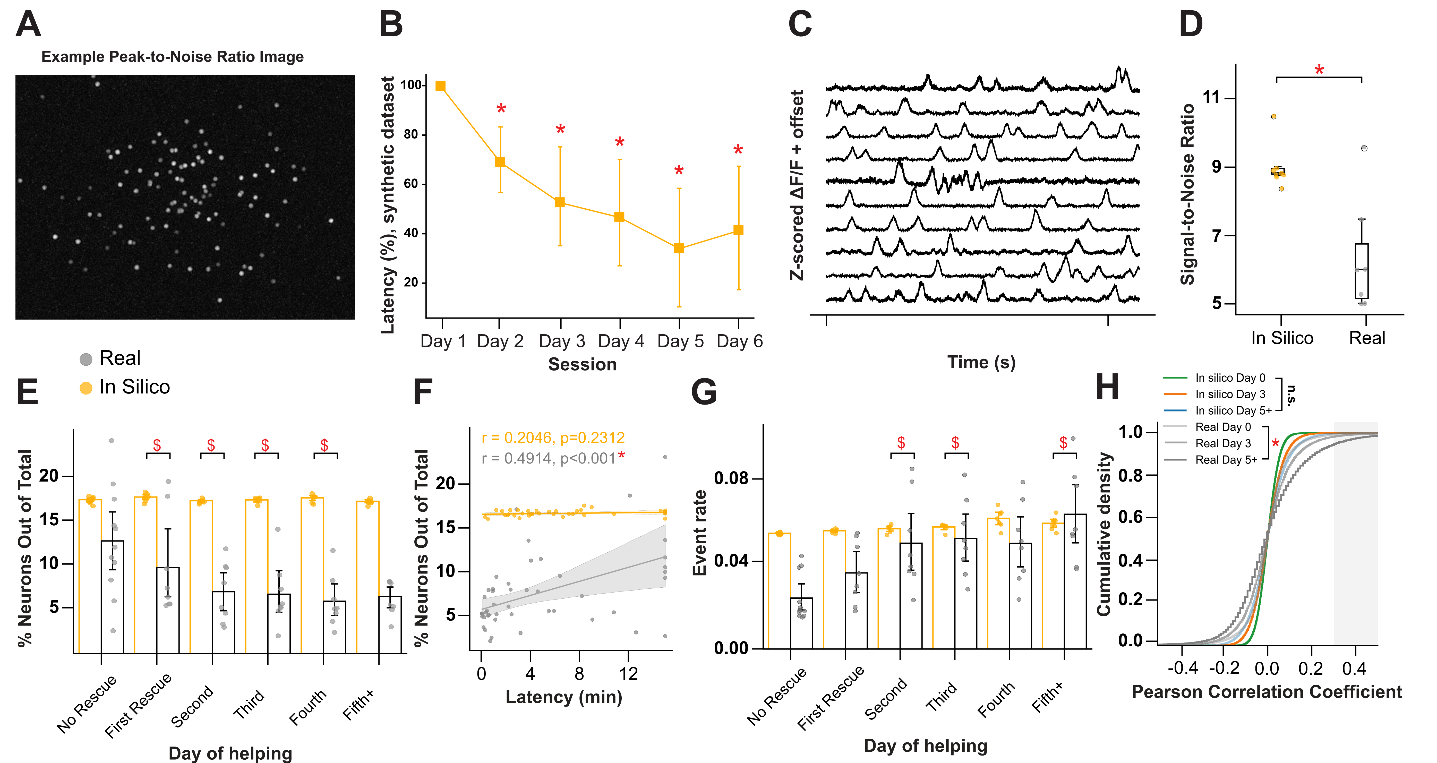

**Figure S3. Validation of neuronal activity analysis using a synthetic calcium imaging dataset, related to Figure 3**. (A) Peak-to-noise image example from one of the simulated recordings. (B) Modeled latency percentages for successful door opening and victim mouse liberation (LMM Omnibus test for day of helping: F(5,35.9) = 26.3, p < 0.001; difference factor coding for day-of-helping comparisons followed by Holm's multiple comparison test). (C) Representative z-scored ΔF/F signals from 10 simulated cells. (D) Box plots displaying the distribution of average signal-to-noise ratios for real (gray) and synthetic (yellow) datasets, with overlaid individual data points representing the mean values of neurons' signal-to-noise ratios from each animal (Mann–Whitney U test: p = 0.014). (E) Comparison of the percentage of neurons in each mouse relative to the total number of neurons across all sessions, grouped by days of helping from 'No Rescue' to '5 or more rescues', between synthetic (yellow) and real (gray) datasets (LMM Omnibus tests: Day of Helping, F(5,80.8)=5.05, p < 0.001; Dataset, F(1,10.8)=87.94, p < 0.001; Day of Helping × Dataset, F(5,80.8)=4.83, p < 0.001). $ indicates days with a significantly different rate of change between datasets (p < 0.05). (F) Pearson correlation between neuron percentage and latency for helping, shown for real (gray) and synthetic (yellow) datasets (r = 0.49, p < 0.001) and synthetic datasets (r = 0.20, p = 0.23). (G) Comparison of the event rate (Hz) per animal across days of helping for real (gray) and synthetic (yellow) datasets (LMM Omnibus tests: Day of Helping, F(5,79.70)=7.94, p < 0.001; Dataset, F(1,8.68)=6.07, p = 0.037; Day of Helping × Dataset, F(5,79.70)=3.67, p = 0.005). $ indicates days with a significantly different rate of change between datasets (p < 0.05). (H) Cumulative distributions of Pearson's coefficients per day of helping; the gray area indicates values of r ≥ 0.3. Bootstrapped two-sample Kolmogorov–Smirnov tests revealed significant differences between Day 0 and Day 5+ for real (gray) dataset (KS = 0.197 ± 0.057 CI, p = 0.048, corrected), while other comparisons remained only marginally significant for other days in real (gray) dataset (Day 0 vs. Day 3: KS = 0.140 ± 0.067 CI, p = 0.585, corrected; Day 3 vs. Day 5+: KS = 0.135 ± 0.057 CI, p = 0.585, corrected) and non-significant for synthetic datasets (Day 0 vs. Day 3: KS = 0.141 ± 0.052 CI, p = 1.0, corrected; Day 0 vs. Day 5+: KS = 0.183 ± 0.070 CI, p = 0.096, corrected; Day 3 vs. Day 5+: KS = 0.113 ± 0.050 CI, p = 1.0, corrected). $ indicates days with a significantly different rate of change between datasets (p < 0.05), * indicates significant difference between groups real an in silico data (p < 0.05). All data are presented as mean ± 95% confidence intervals.

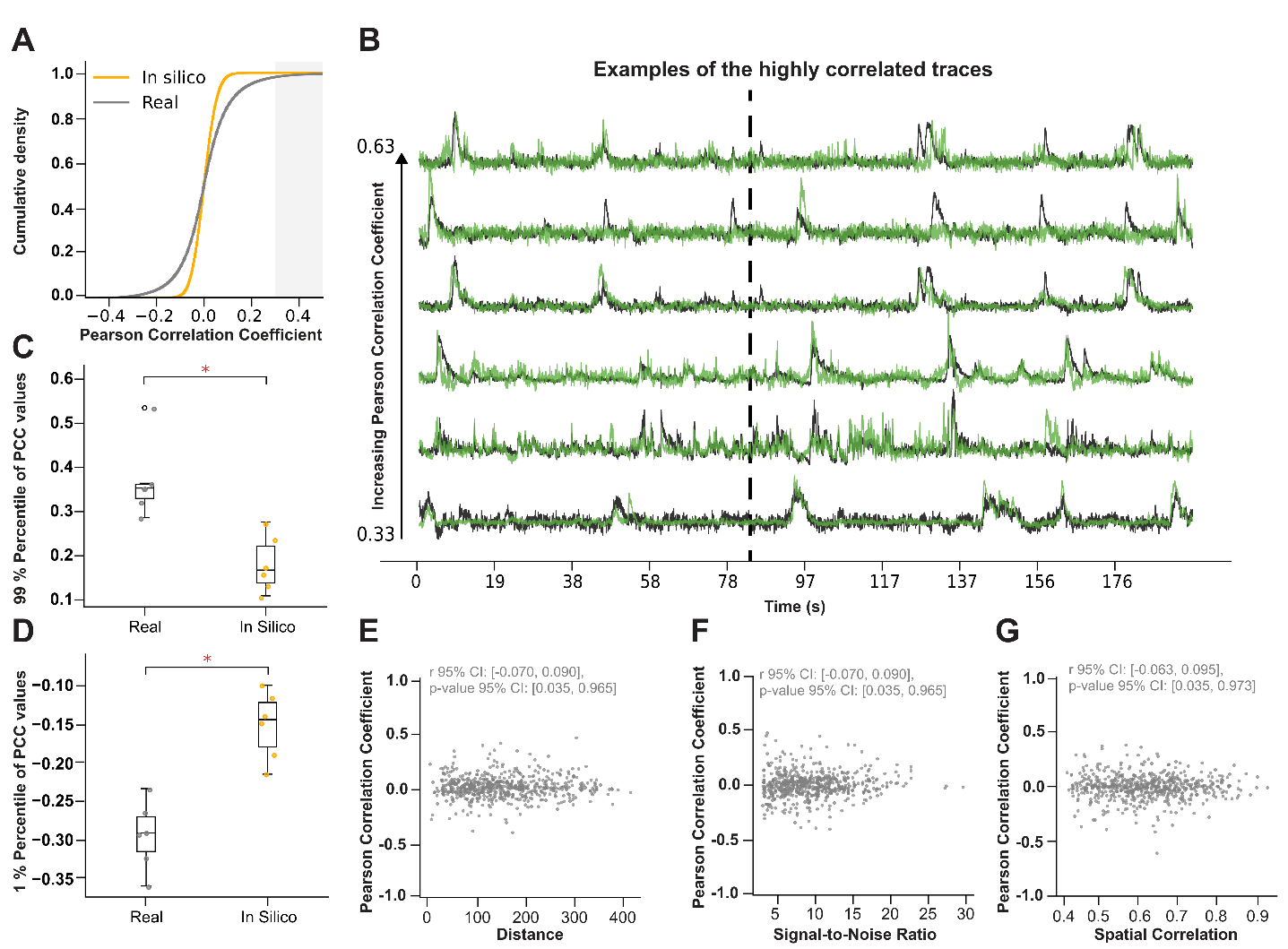

**Figure S4. Validation and evaluation of extrinsic influences on pairwise neuronal correlations, related to Figure 3**. (A) Cumulative distribution function of Pearson correlation coefficients for real (gray) and synthetic (yellow) signals. Pairwise KS tests with bootstrapping yielded a mean KS statistic of 0.138 (95% CI = [0.094, 0.185]) and a corrected p-value of 0.144. (B) Overlaid z-scored ΔF/F signals from highly correlated neuronal pairs (PCC > 0.33) in the real dataset. (C) Box plots showing the 99th percentile of correlation coefficients for real (gray) and synthetic (yellow) datasets, with individual data points representing the mean values from each day of helping (Mann-Whitney U test: p = 0.0016). (D) Box plots showing the 1st percentile of correlation coefficients for real (gray) and synthetic (yellow) datasets, with individual data points representing the mean values from each day of helping (Mann-Whitney U test: p < 0.001). (E) Scatter plot showing the relationship between the distance of neuronal pairs and their correlation coefficients (Pearson r 95% CI: [–0.070, 0.090]; p-value 95% CI: [0.035, 0.965]). (F) Scatter plot showing the relationship between the average signal-to-noise ratio of neuronal pairs and their correlation coefficients (Pearson r 95% CI: [–0.074, 0.073]; p-value 95% CI: [0.065, 0.968]). (G) Scatter plot showing the relationship between the average spatial correlation of neuronal pairs and their correlation coefficients (Pearson r 95% CI: [–0.063, 0.095]; p-value 95% CI: [0.035, 0.973]). For all data presented * indicates group difference with p < 0.05.

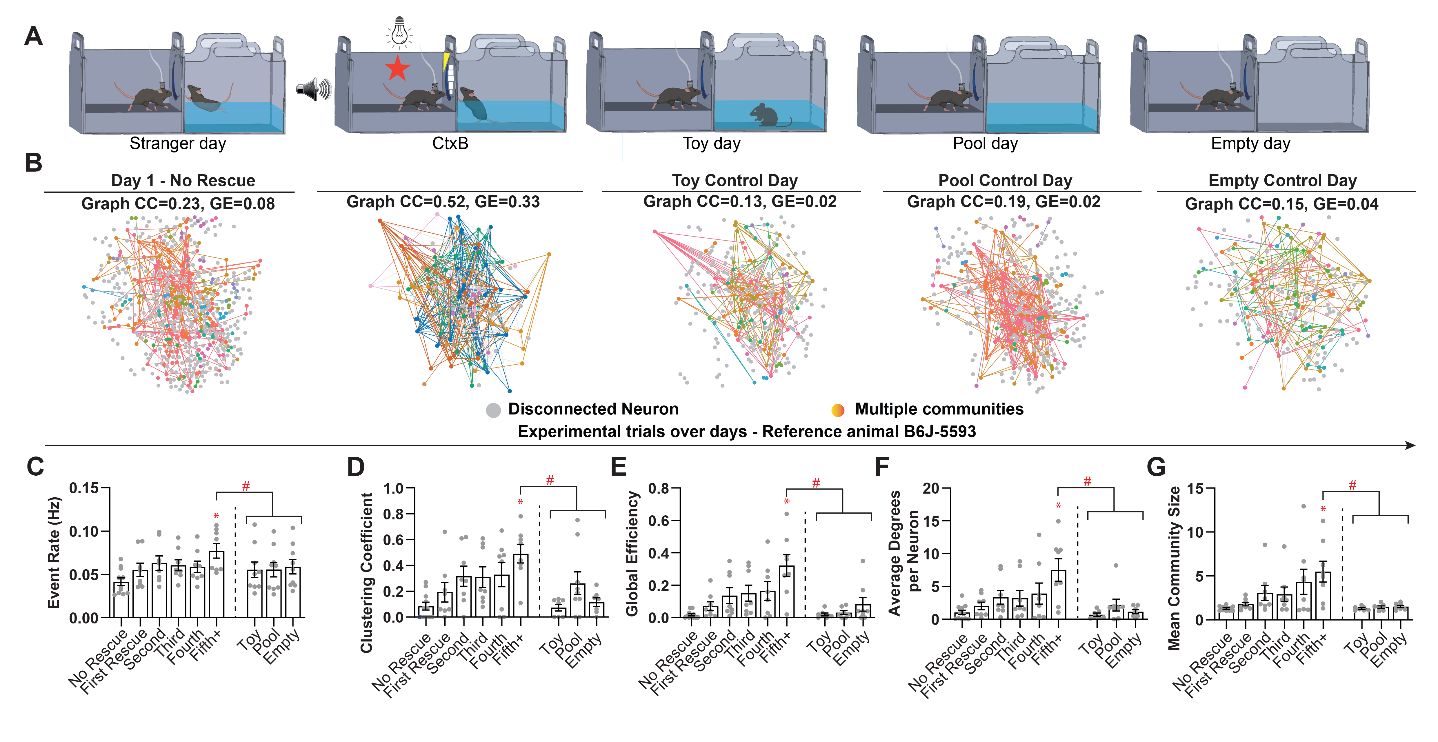

**Figure S5. Dorsal CA1 network is less engaged during control days, related to Figure 3.** (A) Scheme of control days performed after training and recent test sessions. The control days consisted of introducing the trained helper mouse to the same training context with the victim’s chamber having either a toy mouse in the cold water, a pool of cold water without anything else in it or completely empty, respectively. (B) Representative graphs for one animal at the day 1, with no successful rescue (leftmost panel) and day 6 of training, after 5 successful rescues (second from left to right), and on the 3 control days. Each dot is a neuron arranged approximately at the real location in the endoscopic video and each line represents a correlation which coefficient is over +0.30. Gray circles represent disconnected neurons whereas other colored circles represent different unique communities. (C) Event rate (Hz) per animal across days of helping, starting from „No Rescue” to “5 or more rescues” and subsequent control sessions (LMM Omnibus test for Day of Helping: F(8,59) = 6.42, p < 0.0001; difference factor coding to compared with previous levels). (D) Clustering Coefficient across days of rescue (LMM Omnibus test for Day of Helping: F(8,55) = 5.91, p < 0.0001). (E) Global Efficiency of the network across days of helping (LMM Omnibus test for Day of Helping: F(8,54) = 7.59, p < 0.0001). (F) Average degrees per neuron across days of rescue (LMM Omnibus test for Day of Helping: F(8,54) = 4.42, p = 0.0004). (G) Average community size per mouse across days of rescue (LMM Omnibus test for Day of Helping: F(8,53) = 4.17, p = 0.006). All data are presented as mean ± SEM. Outliers were identified and removed using the ROUT method. *p < 0.05 when comparing with all previous rescue trials, #p<0.05 when comparing control sessions with 5+ rescue trials.

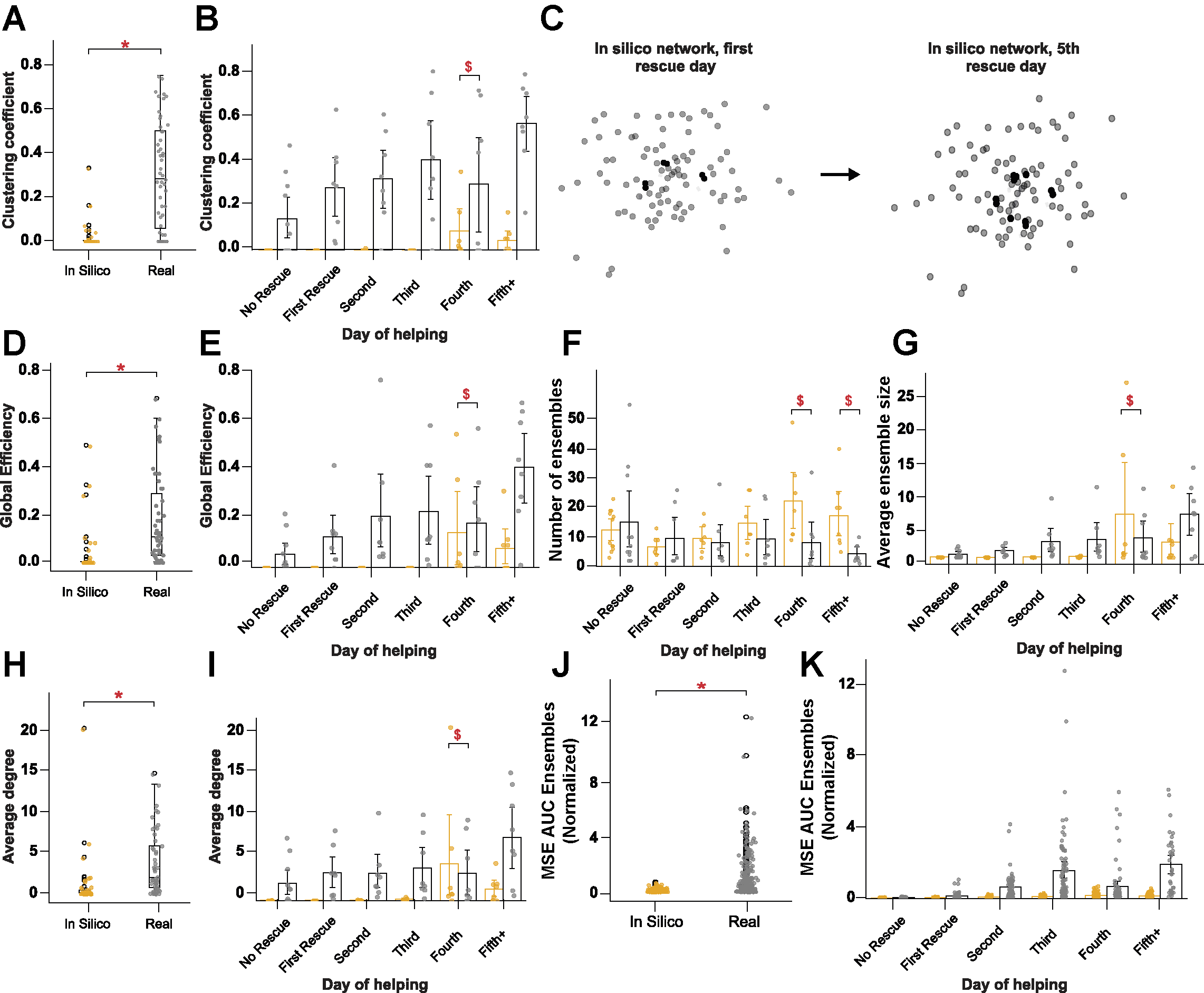

**Figure S6.** **Validation of connectivity analysis using a synthetic calcium imaging dataset, related to Figure 3** (A) Box plots displaying the distribution of clustering coefficients for real (gray) and synthetic (yellow) datasets, where each point represents one animal in one session (Mann–Whitney U test: p < 0.001). (B) Clustering coefficient across days of rescue for real (gray) and synthetic (yellow) datasets (LMM Omnibus tests: Day of Helping, F(5,79.29)=2.15, p = 0.068; Dataset, F(1,8.24)=12.63, p = 0.007; Day of Helping × Dataset, F(5,79.29)=2.38, p = 0.046). $ indicates days with a significantly different rate of change between datasets (p < 0.05). (C) Graphs for the no-rescue day and 5+ days of rescue for one animal from the synthetic dataset. Graphs were constructed using all correlations with coefficients above a threshold of +0.30. (D) Box plots displaying the distribution of global efficiency for real (gray) and synthetic (yellow) datasets, where each point represents one animal in one session (Mann–Whitney U test: p < 0.001). (E) Global efficiency of the network across days of helping for real (gray) and synthetic (yellow) datasets (LMM Omnibus tests: Day of Helping, F(5,79.88)=2.85, p = 0.020; Dataset, F(1,8.97)=6.98, p = 0.027; Day of Helping × Dataset, F(5,79.88)=1.72, p = 0.140). $ indicates days with a significantly different rate of change between datasets (p < 0.05). (F) Number of communities per mouse across days of rescue for real (gray) and synthetic (yellow) datasets (LMM Omnibus tests: Day of Helping, F(5,74.1)=2.01, p=0.087; Dataset, F(1,12.9)=2.06, p=0.175; Day of Helping × Dataset, F(5,74.1)=4.75, p<0.001). $ indicates days with a significantly different rate of change between datasets (p < 0.05). (G) Average community size per mouse across days of rescue for real (gray) and synthetic (yellow) datasets (LMM Omnibus tests: Day of Helping, F(5,74.3)=4.67, p < 0.001; Dataset, F(1,14.2)=1.30, p = 0.274; Day of Helping × Dataset, F(5,74.3)=2.07, p = 0.079). $ indicates days with a significantly different rate of change between datasets (p < 0.05). (H) Box plots displaying the distribution of average degree per neuron for real (gray) and synthetic (yellow) datasets, where each point represents one animal in one session (Mann–Whitney U test: p < 0.001). (I) Average degrees of neurons across days of rescue for real (gray) and synthetic (yellow) datasets (LMM Omnibus tests: Day of Helping, F(5,73.85)=2.30, p = 0.054; Dataset, F(1,9.69)=4.65, p = 0.057; Day of Helping × Dataset, F(5,73.85)=2.56, p = 0.035). $ indicates days with a significantly different rate of change between datasets (p < 0.05). (J) Box plots displaying the distribution of Multi-Sample Entropy area under curve values (normalized by latency to rescue) for real (gray) and synthetic (yellow) datasets, where each point represents one community (Mann–Whitney U test: p < 0.001). (K) Multi-Sample Entropy area under curve value (normalized by latency to rescue) per community across test sessions for real (gray) and synthetic (yellow) datasets (LMM Omnibus tests: Day of Helping, F(5,11.0)=14.54, p<0.001; Dataset, F(1,12.7)=13.22, p=0.003; Day of Helping × Dataset, F(5,11.0)=8.85, p=0.001). $ indicates days with a significantly different rate of change between datasets (p < 0.05). , * indicates significant difference between groups real an in silico data (p < 0.05). All data are presented as mean ± 95% confidence intervals. n.s. = non-significant.

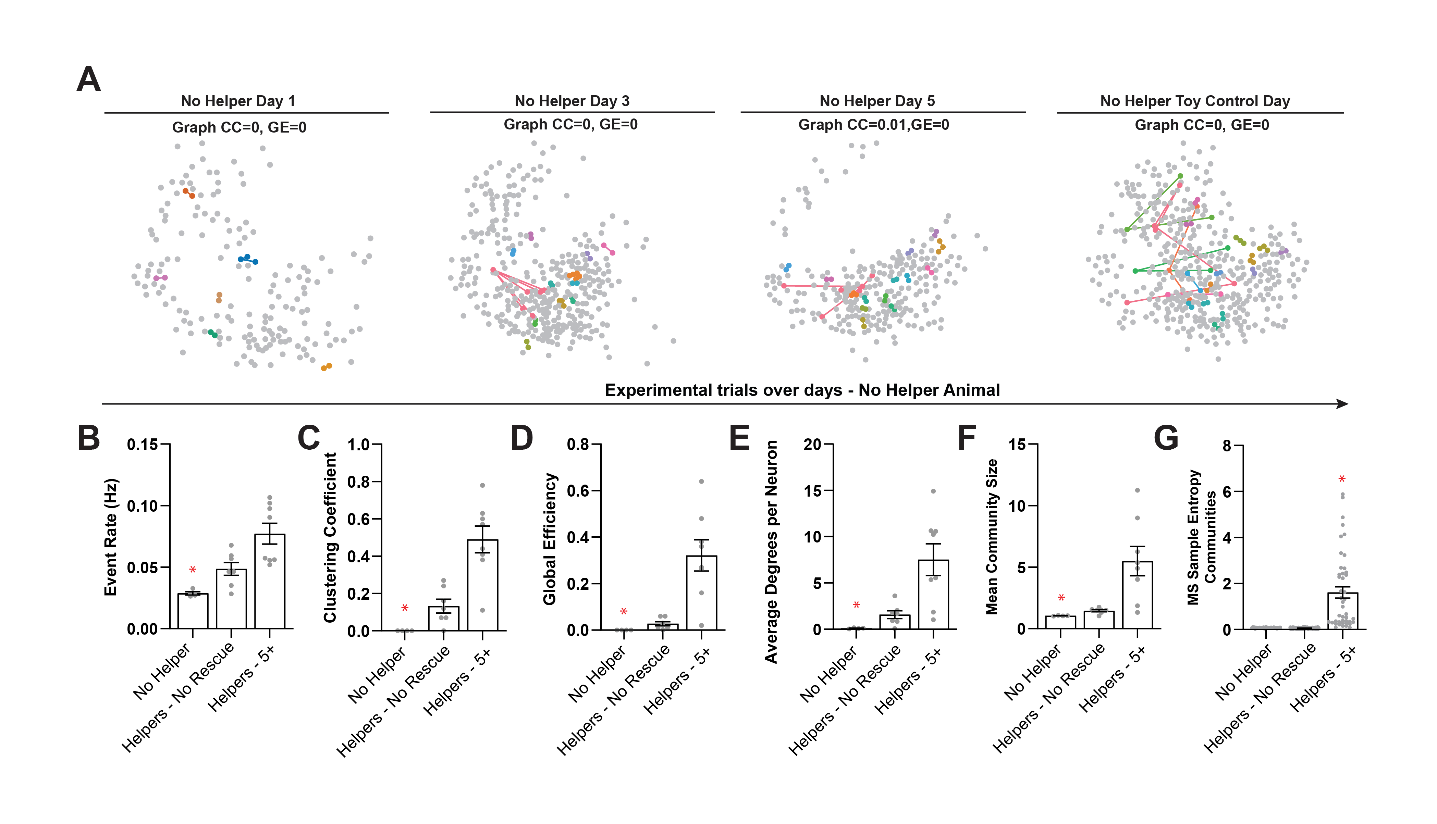

**Figure S7. Engagement of dorsal CA1 for the non-helper mouse did not increase over days, related to Figure 3.** (A) Representative graphs for the non-helper animal at the day 1 (leftmost panel), day 3 (second from left to right) and day 5 (third from left to right) of training, and on one of the 3 victimless control days. Each dot is a neuron arranged approximately at the real location in the endoscopic video and each line represents a correlation which coefficient is over +0.30. Gray circles represent disconnected neurons whereas other colored circles represent different unique communities. (B) Event rate (Hz) per animal across days of helping, comparing the No-helper animal to the „No Rescue” and “5 or more rescues” sessions of the helper animals (Kruskal-Wallis: KW = 10.60, p = 0.0012). (C) Clustering Coefficient of the No-helper compared to helpers (Kruskal-Wallis: KW = 13.40, p < 0.0001). (D) Global Efficiency of the network of the No-helper compared to helpers (Kruskal-Wallis: KW = 13.64, p < 0.0001). (E) Average degrees per neuron of the No-helper compared to helpers (Kruskal-Wallis: KW = 12.18, p = 0.0002). (F) Average community size per mouse of the No-helper compared to helpers (Kruskal-Wallis: KW = 12.18, p = 0.0012). (G) Normalized MSE area under curve value per community of the No-helper compared to helpers: (Kruskal-Wallis: KW = 122.5, p < 0.0001). All data are presented as mean ± SEM. n.s. = non-significant, *p < 0.05 when comparing with helpers no rescue and 5+.
